# Broadly Reactive Anti-VHH Antibodies for Characterizing, Blocking, or Activating Nanobody-Based CAR-T Cells

**DOI:** 10.1101/2024.09.18.613561

**Authors:** Scott McComb, Bianca Dupont, Alex Shepherd, Bigitha Bennychen, Anne Marcil, Mehdi Arbabi-Ghahroudi, Laura Tamblyn, Shalini Raphael, Joey Sheff, Greg Hussack, Anna N. Moraitis, Cunle Wu, Annie Aubry, Christine Gadoury, Julie Lippens, Martine Pagé, Annie Fortin, Simon Joubert, Linda Lamoureux, Marie Parat, Pierre Plante, Félix Malenfant, Mauro Acchione, Petra Pohankova, Joe Schrag, Andrea Acel, Mathieu Coutu, Emma Smith, Majida El Bakkouri, Jennifer J. Hill, Tammy-Lynn Tremblay, Manceur Aziza, Sharlene Faulkes, John Webb, Ahmed Zafer, Qin Zhu, Tina Nguyen, Robert A. Pon, Risini D. Weeratna

## Abstract

Production of chimeric antigen receptor T cell (CAR-T) therapies is dependent on the use of antibody reagents to label, isolate, and/or expand T cell products. We sought to create antibody-based tools that directly target the variable domain of heavy-chain only antibodies (VHH or nanobody) used in some CAR molecules. Two murine antibodies were identified which bind to distinct epitopes in the conserved framework regions of llama-derived VHHs, and not to human VH domains. We produced a high-quality dual-clonal anti-VHH antibody product which reacts with over 98% of VHH proteins, regardless of their antigenic specificity. Anti-VHH binding did not disrupt VHH/antigen interaction, and thus could be used for secondary labeling to assess cellular or tissue reactivity of VHH molecules. Despite not interfering with antigen binding, anti-VHH antibodies potently inhibited VHH-CAR function, blocking CAR-T activation and cytolytic killing of target cells. When immobilized, anti-VHH antibodies could also be applied for activation and expansion of VHH CAR-T cells, inducing 730-fold mean expansion, >94% CAR purity, with retained CD8/CD4 heterogeneity. Functionally, anti-VHH antibody-expanded CAR-T cells maintained strong antigen specific activity without functional exhaustion. Overall, these data identify a useful new tool for understanding and manipulating VHH-based CAR-T cells.

**Funding Source:** This work was funded by the National Research Council Canada Disruptive Technology Solutions Cell and Gene Therapy challenge program, and BioCanRx

**Declaration of interests:** The anti-VHH antibodies reported here are the subject of a provisional patent application by the National Research Council of Canada

## Introduction

The development of CD3-targeting antibodies (Abs) was a pivotal step towards creating a standardized method for expanding T-cells, eventually leading to the development of modern T-cell therapies such as chimeric antigen receptor T-cell therapy. As with other receptor targeting antibodies, soluble CD3-specific Abs can block antigen-specific stimulation of T-cells through the TCR, whereas when these same anti-CD3 Abs are multimerized they will act as potent activators of T-cell proliferation^1,2^. Cross-linking of anti-CD3-Abs, either through Fc-conjugation^3^, surface absorption^4^, multimerization^5,6^, artificial antigen presenting cells^7^, or bead conjugation^8^, are all proven strategies to exploit the T-cell activating properties of anti-CD3-Abs. In addition to altering T-cell receptor signaling, anti-CD3 Abs are among a plethora of Abs that are directly conjugated to chemical fluorophores for identification of T cells via flow cytometry and/or cell isolation via magnetic or flow-based cell sorting.

Given the indispensability of CD3-targeted Abs to the scientific and clinical development of T cell therapies, we reasoned that establishing a standardized Ab-based reagent targeting the antibody framework domains within the antigen-binding domain of CAR-T receptors could have similar multifaceted utility for the development of CAR-T therapies. Specifically, our focus has been on the development of CAR therapies utilizing variable domains of llama heavy-chain only antibodies (VHHs), which are also known as nanobodies. VHHs have several advantages over single-chain variable fragment domains (scFvs) which are more widely used in CAR therapies, including a simpler single-gene structure, smaller size, and higher genetic homology to human variable domains than mouse or rabbit antibodies^9^. We have recently reported on novel VHH-based CAR molecular discovery and optimization for VHH-CARs targeting EGFR and CD22^10,11^. The most clinically advanced application of VHH binding domains in a CAR is in ciltacabtagene autoleucel, a BCMA-targeted CAR-T product now approved as a marketed treatment of multiple myeloma in a growing number of countries^12^.

Here we sought to identify murine monoclonal antibodies (mAbs) that can selectively bind to llama-derived VHHs of various VHH-CAR proteins, without cross-reacting with human or mouse-derived Abs, and explore their potential applications. We produced a high quality and consistent mAb product that can be used as a secondary anti-VHH staining reagent to characterize VHH binding to antigen-expressing target cells, or to quantify surface expression of VHH-CARs. These mAbs show broad reactivity to almost all VHHs tested, regardless of their antigenic specificity. In addition to staining applications, we also find that anti-VHH mAbs can recapitulate other functions of CD3-specific Abs such as blockade of antigen specific response or activating specific expansion of VHH CAR-T cells.

## Results

### Identification of broadly cross-reactive anti-VHH mAbs

In order to generate mAbs specific for VHH domains, a previously reported blood-brain barrier transmigrating VHH (clone FC5)^13^ was produced and purified from bacterial culture via affinity column purification. Four A/J mice were then immunized with VHH emulsified in TiterMax adjuvant, and boosted with VHH before harvesting spleens. Hybridoma cultures were then generated via electrofusion and clone screening for VHH binding as previously reported^14^. A total of 233 hybridoma supernatants were screened revealing 37 supernatants with reactivity to FC5. To avoid potential problematic reactivity with human Abs in downstream application, 31 of 37 hybridoma supernatants were specifically selected for negative reactivity via ELISA against polyclonal human IgG containing serum (Figure 1A, Supplemental Table 1). Of the 31 remaining hybridoma supernatants, only 10 showed broad reactivity to 3 llama VHHs tested via ELISA. Finally, 7 supernatants were further selected based on hybridoma growth and antibody productivity characteristics.

**Figure 1:**
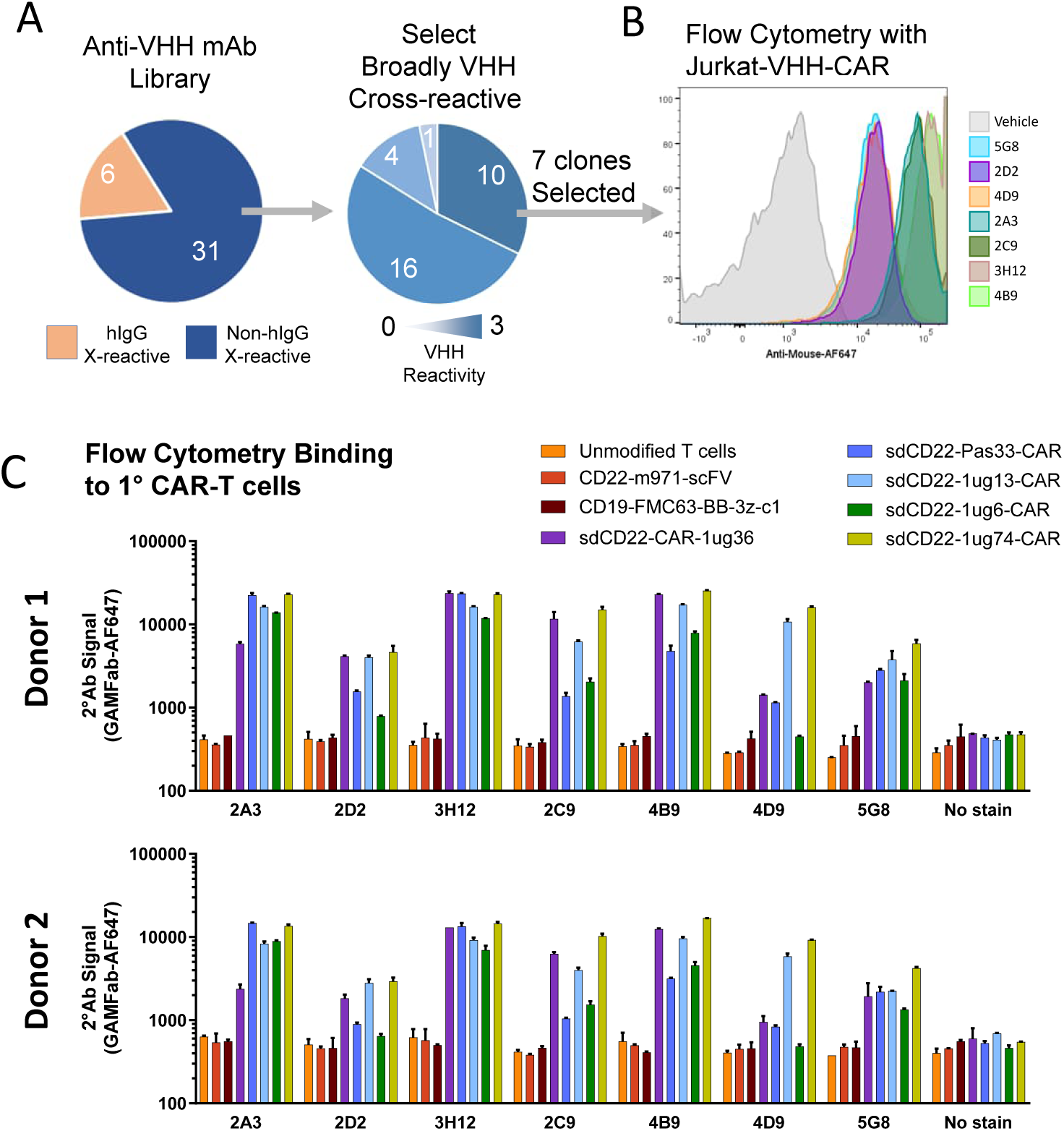
Selection of 2 broadly cross-reactive mouse anti-VHH mAbs A mouse was immunized with a purified VHH, B cells were collected and subjected to hybridoma electrofusion. (A) Clonal hybridoma culture supernatant were then screened via ELISA for reactivity to human total IgG protein, selecting only those with negative reactivity to human IgG for further testing. Non-human reactive supernatants were then screened via ELISA for cross reactive binding to 3 different VHHs, with a total of 10 clones showing reactivity to 3 of 3 VHHs. Flow cytometry was then conducted with Jurkat cells bearing stable expression of surface linked VHH-chimeric antigen receptors (VHH-CAR). (B) Seven anti-VHH hybridoma supernatants were then screened via secondary staining fluorescent anti-mouse IgG and flow cytometry to examine reactivity against primary human T cells bearing 5 divergent VHH-CAR proteins or scFv-based CARs. (C) Anti-VHH hybridoma supernatants were similarly screened for reactivity against primary CAR-T cells from 2 different healthy donor blood samples expressing 5 different VHH-CARs, 2 control scFv-CARs, or untransduced T cells. Bar graphs show the mean of one experiment performed in duplicate +/− SEM. Supernatants with the broadest cross-reactivity (2A3 and 3H12) were selected for downstream testing.

As our primary intent for generating anti-VHH mAbs was for the surface labeling of VHH-CAR expressing cells, we first confirmed via flow cytometry reactivity of all 7 candidate supernatants to Jurkat cells stably expressing a VHH-CAR (Figure 1B). Next, we performed the same analysis of surface reactivity against 5 divergent VHH-CAR molecules expressed in lentivirally-transduced primary human T cells derived from two healthy donors, generated as reported^15^. Results revealed varying reactivity to VHH-CAR expressing cells, with two particular mAb supernatants (clones 2A3 and 3H12) showing strong and consistent reactivity across all 5 VHH-CARs tested (Figure 1C). Neither anti-VHH mAb supernatant showed reactivity against control murine and human scFv-CARs, FMC63 and m971, or against unmodified mock transduced T cells. Overall, these results identified two promising candidate anti-VHH mAbs with broad reactivity to VHH domains.

### Production and testing of recombinant anti-VHH mAbs

Having selected two candidate anti-VHH mAbs, the heavy and light chain variable genes were sequenced from 2A3 and 3H12 mAb IgG1 hybridoma clones. The variable domains from these clones were then recombined to mouse IgG2a-Fc backbones. It has been our experience that murine IgG2a antibodies result in higher antibody production and have better stability. Binding to purified VHH shows that recombinant IgG2a anti-VHH (rec2A3 and rec3H12) mAbs maintain similar reactivity to VHH as the original IgG1 mAbs (Figure 2A). We also confirmed that the recombinant IgG2a anti-VHH mAbs maintain low reactivity to human IgG (Supplemental Figure 1A). As product quality and consistency is vital for use in downstream pre-clinical and clinical processes of CAR-T, we also confirmed inter-batch consistency, freeze-thaw stability, and stability over at least 12 months in −80°C storage for this product (Supplemental Figure 1B-F). Given these strong product quality and consistency observations, we used the recombinant IgG2a forms of anti-VHH mAbs for further characterization and application studies discussed throughout the remainder of this manuscript.

**Figure 2:**
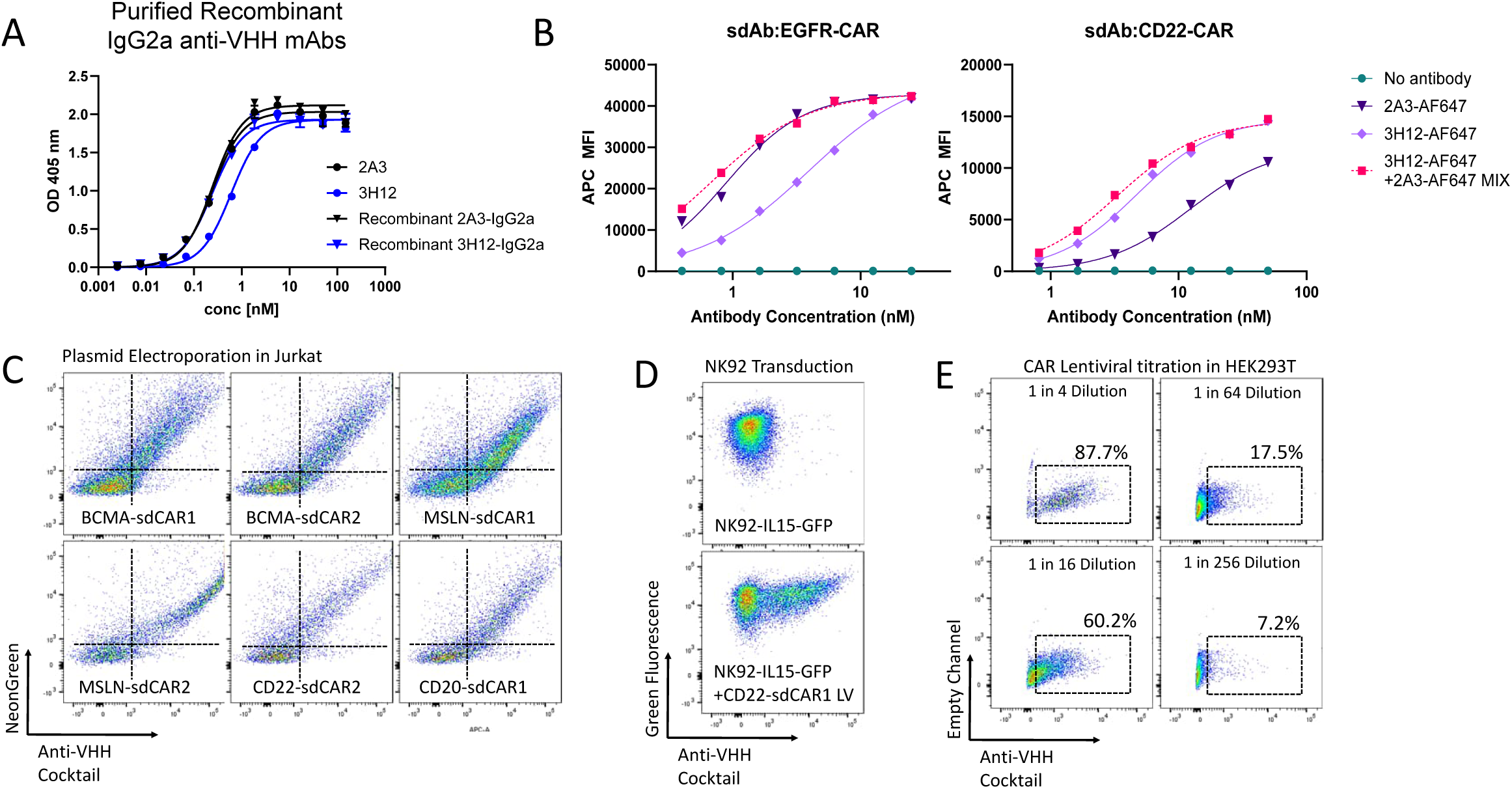
A mixed antibody product of recombinant anti-VHH-IgG2a antibodies can be applied to quantitate surface VHH-CAR expression. The heavy and light chain variable genes were sequenced from selected 2A3 and 3H12 hybridoma clones. The variable domains from these clones were then fused to mouse IgG2a backbones to improve productivity and stability of purified antibodies. (A) ELISA was performed against purified VHH using purified IgG1 anti-VHH mAbs or recombinant IgG2a anti-VHH mAbs (rec2A3 and rec3H12). (B) Recombinant anti-VHH mAbs were directly labelled with AlexaFluor-647 and used individually or as a 1:1 molar ratio mix of both antibodies. VHH-CAR Jurkat cells known to have lower affinity for either 3H12 (EGFR-CAR, *left*) or 2A3 (CD22-CAR, *right*) were stained and examined via flow cytometry. (C) VHH-CAR expressing cells were generated expressing 6 divergent CARs, stained with the AF647 anti-VHH antibody cocktail and assessed via flow cytometry. (D) Similarly, VHH-CAR expression on stable IL15-GFP expressing natural killer (NK92-IL15) cells was also confirmed using anti-VHH staining. (E) HEK293 cells were used to titrate non-fluorescently labelled VHH-CAR lentiviral particles by using varying dilutions of viral supernatant on HEK cells via AF647 anti-VHH staining and flow cytometry.

Recombinant anti-VHH mAbs were then directly labelled with AlexaFluor-647 and used in various fluorescent flow cytometric analyses of VHH-CAR expressing cells. Importantly, early testing revealed that certain VHH-CAR molecules were not labelled efficiently with 3H12 (EGFR-CAR, Figure 2B *left*) or 2A3 (CD22-CAR, Figure 2B *right*) respectively. We hypothesized that a 1:1 mix of both anti-VHH mAbs would provide a more consistent staining reagent that could be applied across varying VHH-CAR molecules. Indeed, titration of a mixed anti-VHH product revealed a uniform staining profile in the two VHH-CARs tested here (Figure 2B, broken lines).

To confirm consistent reactivity of this anti-VHH mAb mixture against more VHH-CAR molecules, 6 additional VHH-CAR expressing Jurkat T cell lines with specificity for 4 distinct surface targets (BCMA, mesothelin, CD22, CD20) were stained and analyzed by flow cytometry. Across the 6 additional VHH-CARs, staining with the anti-VHH mAb mix correlated directly with the NeonGreen fluorescent marker co-expressed with VHH-CAR genes (Figure 2C). Similarly, we were also able to use the anti-VHH mAb mix to confirm CAR expression in NK92-IL15 cells transduced with a VHH-CAR lentivirus, wherein stable GFP expression precluded the use of NeonGreen to quantitate CAR expression (Figure 2D). A similar application of anti-VHH mAbs would be to perform a rapid quantification of lentiviral particle production for a VHH-CAR lentivirus without the need to express a fluorescent marker gene (Figure 2E). Overall, these results indicate that a mixed product of anti-VHH mAbs can be used to quantify surface expression of VHH-CAR molecules with divergent variable domain sequences and antigen specificity.

### Anti-VHH mAbs Show Differential Reactivity with Purified VHH Proteins

Next, we wanted to confirm the breadth of reactivity with a large panel of VHHs. We performed an enzyme-linked immunosorbent assay (ELISA), wherein a wide array of VHHs generated over 8 independent VHH discovery projects targeting 11 unique antigens were tested for binding with our anti-VHH mAbs. VHHs were absorbed onto ELISA plates and probed with 3H12, 2A3, or a 1:1 mix of both mAbs, detecting anti-VHH mAb reactivity with an anti-mouse secondary Ab. We found that 75 of 76 unique VHHs are recognized by a the 1:1 mix of anti-VHH mAbs; 74 of 76 are recognized by 3H12 mAb; and 73 of 76 are recognized by 2A3 mAb (Figure 3A). Both mAbs were also able to recognize a recently reported llama VH domain^16^, which was monomeric but distinct in sequence from a typical VHH (Figure 3B). Importantly, we did observe examples wherein 3H12 or 2A3 each showed greater reactivity, such as VHH-59 or VHH-60 respectively, with the mix being the most broadly reactive (Figure 3A,C). Surface plasmon resonance (SPR) assays, using VHHs as analytes flowed over anti-mouse Fc captured 3H12 and 2A3 mAbs, revealed a similar pattern of reactivity to the ELISAs for a representative set of VHHs (Figure 3D), with monovalent binding affinities ranging from *K*_D_ = 3.9 – 9.5 nM and *K*_D_ = 2.4 – 192 nM for 3H12 and 2A3, respectively.

**Figure 3:**
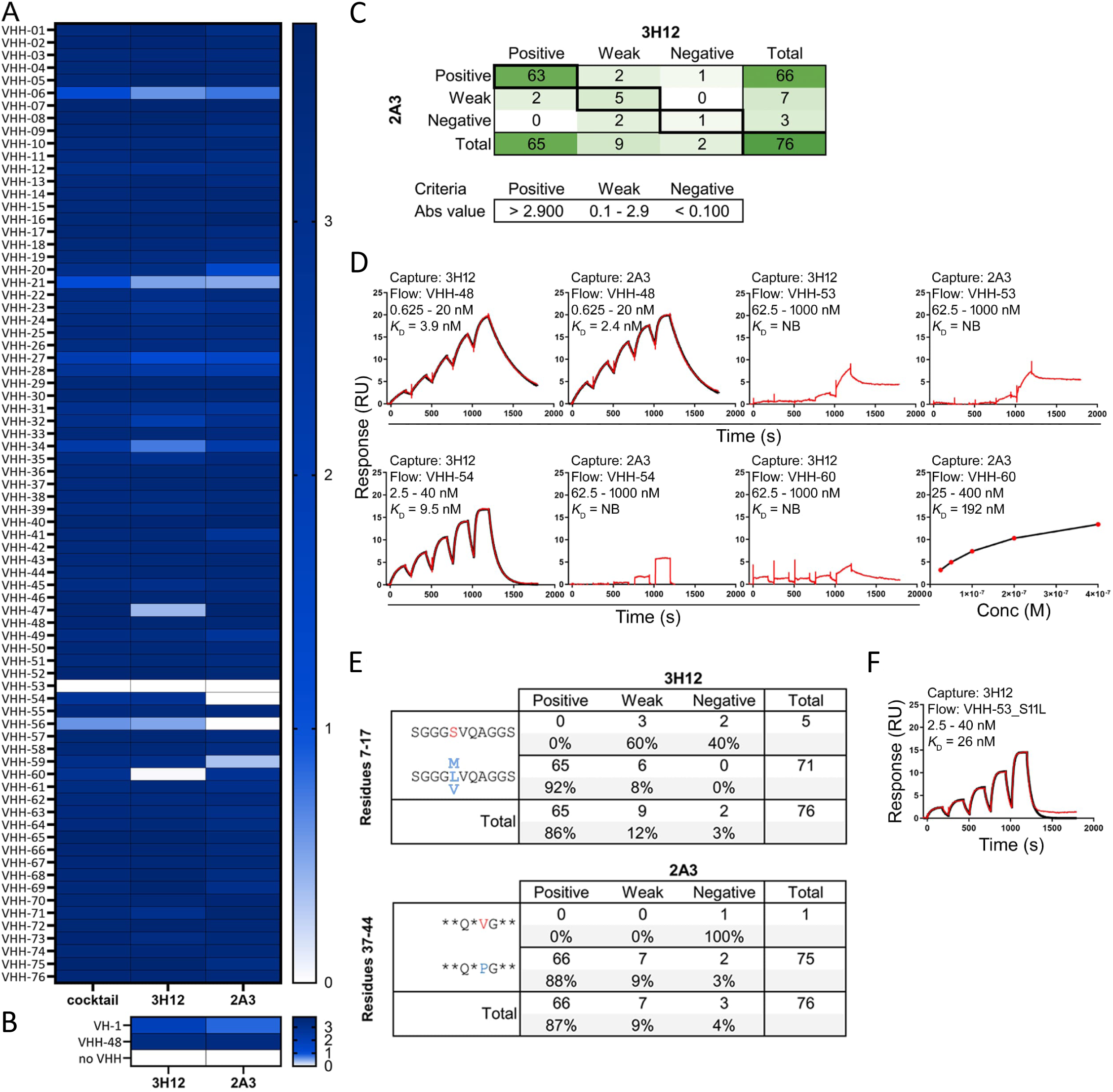
ELISA and SPR-based reactivity of 3H12 and 2A3 mAbs. (A) ELISA demonstrating 3H12 and 2A3 reactivity towards 76 passively-absorbed llama-derived VHHs. “Cocktail” refers to an equimolar mix of 3H12 and 2A3. Binding of mAbs to VHHs was detected with donkey-anti-mouse IgG-HRP. Values are absorbance readings at 450 nm. (B) Similarly, reactivity of 3H12 and 2A3 to a Llama-VH domain, or a control VHH domain were tested via ELISA, (C) Summary of VHH detection by 3H12 and 2A3. (D) SPR sensorgrams showing monovalent binding affinities of select VHHs for mAbs. 3H12 and 2A3 mAbs were captured and VHHs flowed at the concentration ranges indicated. NB: no binding. (E) VHH consensus sequences in FR1 (3H12) and FR2 (2A3) that may be predictors of mAb reactivity. (F) SPR sensorgram showing a single S11L mutation in FR1 of VHH-53 imparts 3H12 binding.

Following these results, we performed a sequence alignment of the 76 VHHs used in ELISA to identify potential sequence determinants associated with positive or negative anti-VHH mAb reactivity. For 3H12, we observed weak or negative reactivity with 5 of 5 VHHs with a serine substitution in position 11 in framework region 1 (FR1) of the VHH (Figure 3E). Introduction of an S11L mutation into VHH-53 was able to restore 3H12 binding (Figure 3F), pinpointing residue 11 as a key interaction for 3H12. For 2A3, we were unable to identify any consensus framework region sequences that were associated with positive or negative binding; however, a motif in FR2 was identified as a potential region important for 2A3 binding (Figure 3E). Overall, these data indicate that the anti-VHH mAbs generated here show broad reactivity with llama VHHs, likely through recognition of distinct epitopes in the framework domains of llama VHHs.

### Hydrogen Deuterium Exchange Mapping of Anti-VHH mAbs Mapping Reveals Distinct but Competitive Epitopes

To further investigate how the anti-VHH mAbs interact with VHHs, we generated a yeast surface display (YSD) library with progressively truncated VHHs, probing with anti-VHH mAbs via whole-cell ELISA. Consistent with our observations in the VHH ELISA above, both 2A3 and 3H12 showed strong binding in a whole-cell ELISA using yeast displaying a cell-wall linked VHH protein. Binding with 3H12 was completely lost with an N-terminal 15 amino acid truncation of the VHH FR1 (Figure 4A, B), but 2A3 maintained partial reactivity. Both mAbs showed negative reactivity to a 21 amino acid or greater truncation. C-terminal truncations also showed negative reactivity but we hypothesize that many of these truncations likely lead to severe misfolding and/or instability of the VHH in YSD. Nonetheless, these data provide supporting evidence that 2A3 and 3H12 show differential binding and likely target distinct epitopes.

**Figure 4:**
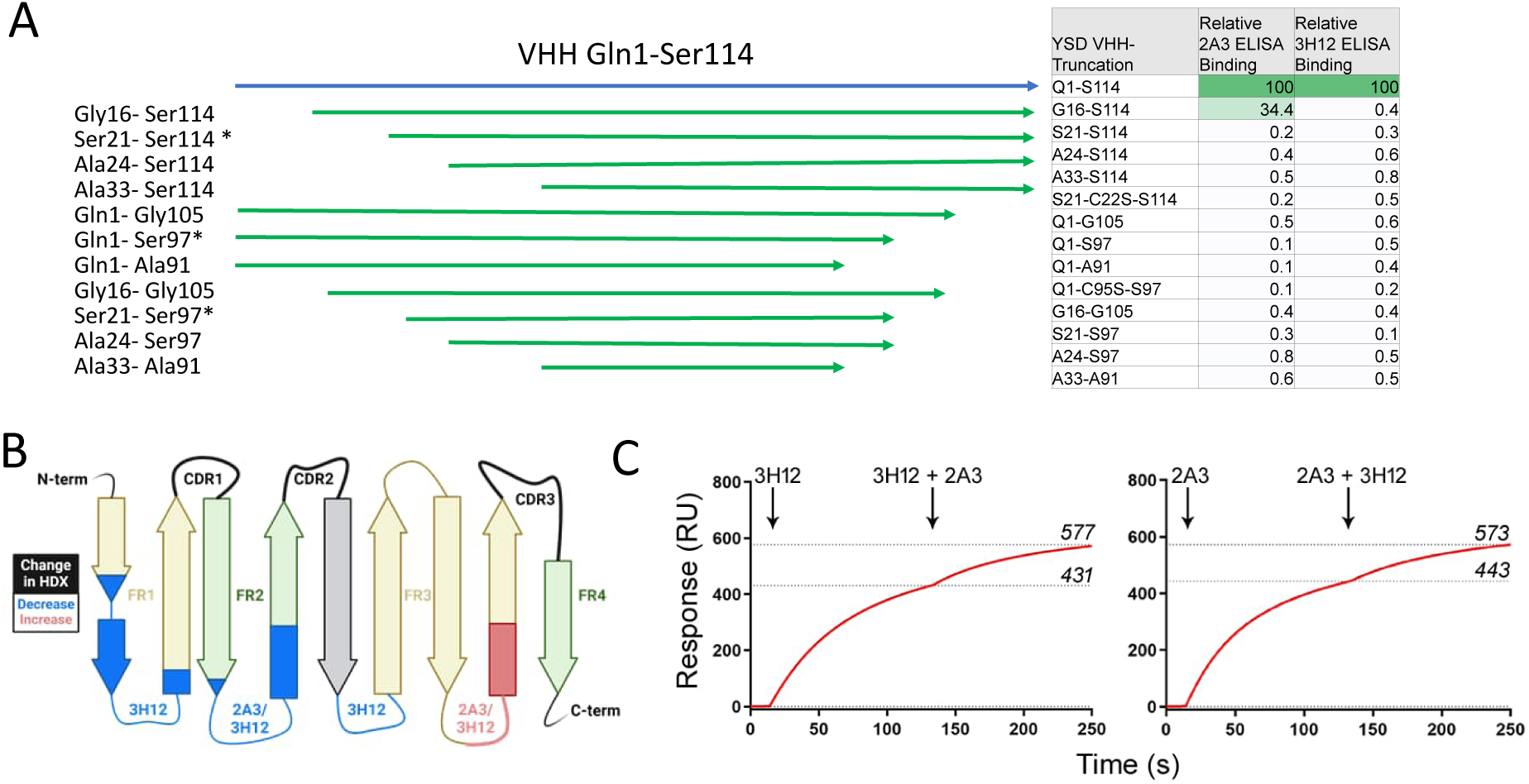
Anti-VHH mAbs 3H12 and 2A3 bind to unique but adjacent epitopes (A) Yeast surface display was used to map binding. Inset table shows the % binding of anti-VHH mAbs via yeast whole-cell ELISA, normalized to the full length VHH. (C) Hydrogen deuterium exchange mass spectrometry was performed on purified VHH alone or bound to excess anti-VHH mAbs. The VHH domain-wire diagram shows regions of reduced (*blue*) or increased (*red*) hydrogen exchange when a VHH test material was saturated with 2A3 or 3H12 anti-VHH mAb. (C) Sensorgrams showing SPR-based competition assays. A VHH was amine coupled and mAbs were sequentially injected. Binding of the first mAb partially prevented binding of the second mAb, regardless of injection order. Response levels (RUs) are shown at the end of each injection.

To more finely map the anti-VHH mAb epitopes, we employed hydrogen deuterium exchange mass spectrometry (HDX-MS). Deuteration was measured for 41 peptides covering >80% of the CD22-specific VHH (clone 1ug36)^10^ test material (Supplemental Figure 2). Results show that saturation of the VHH with 3H12 but not 2A3 resulted in a slower exchange of hydrogen atoms mostly within the N-terminal amino acids of FR1 of the VHH (residues S6-S17), corroborating our YSD findings (Figure 4B, Supplemental Table 2). Additionally, 3H12 slowed HDX for residues within FR2 (Q39-E46) and FR3 (V63-F67), suggesting a discontinuous epitope spanning multiple framework loops that are distal from the hypervariable complementarity determining regions (CDRs). Saturation of the VHH protein with 2A3 mAb only resulted in slower HDX within the FR2 region (Figure 4B). Both VHH proteins exhibited faster exchange in residues D89-Y94 (red, Figure 4B) which likely results from allosteric conformational changes induced by stabilization of the framework loops.

Competitive SPR binding experiments involving sequential anti-VHH mAb injections over an amine-coupled VHH surface revealed 2A3 and 3H12 partially competed with one another, with binding of the first mAb preventing full unobstructed binding of the second mAb (Figure 4C). Given the much larger size of the mAbs compared to a VHH (∼10× larger), the inability of the anti-VHH mAbs to simultaneously bind with a VHH is unsurprising. Collectively our sequence analyses, ELISA, YSD, HDX-MS, and SPR results, indicate that the anti-VHH mAbs recognize two distinct but adjacent epitopes within the conserved FR domains of VHHs, which are distal from the CDR.

### Anti-VHH mAbs for Assessment of VHH Cellular Reactivity

Based on our epitope mapping experiments, we expected that anti-VHH mAb binding should not disrupt CDR-mediated VHH-antigen interaction, thus we wanted to confirm whether these mAbs could be applied to investigate cellular reactivity of soluble VHH proteins. Antigen expressing or deficient cells were stained with varying concentrations of VHHs, washed, and then probed with a fluorophore conjugated anti-VHH mAb mix. Flow cytometry assessment showed specific reaction of anti-CD22 VHHs with CD22 expressing cells but not CD22-knockout cells (Figure 5A). Similarly, BCMA-specific VHHs showed reactivity to a human multiple myeloma cell line, but not against BCMA-negative human Jurkat T cells (Figure 5B). To see whether a similar technique could be applied to examine the tissue distribution of VHH binding, we also tested an immunohistochemistry staining method using an anti-mouse-HRP polymer tertiary antibody detection reagent (Figure 5C). Using this method, human tonsillar samples stained with CD22-specific VHHs revealed characteristic follicular staining pattern consistent with human B cells. These results demonstrate that anti-VHH mAbs are effective for characterization of cell and tissue binding of novel VHHs.

**Figure 5:**
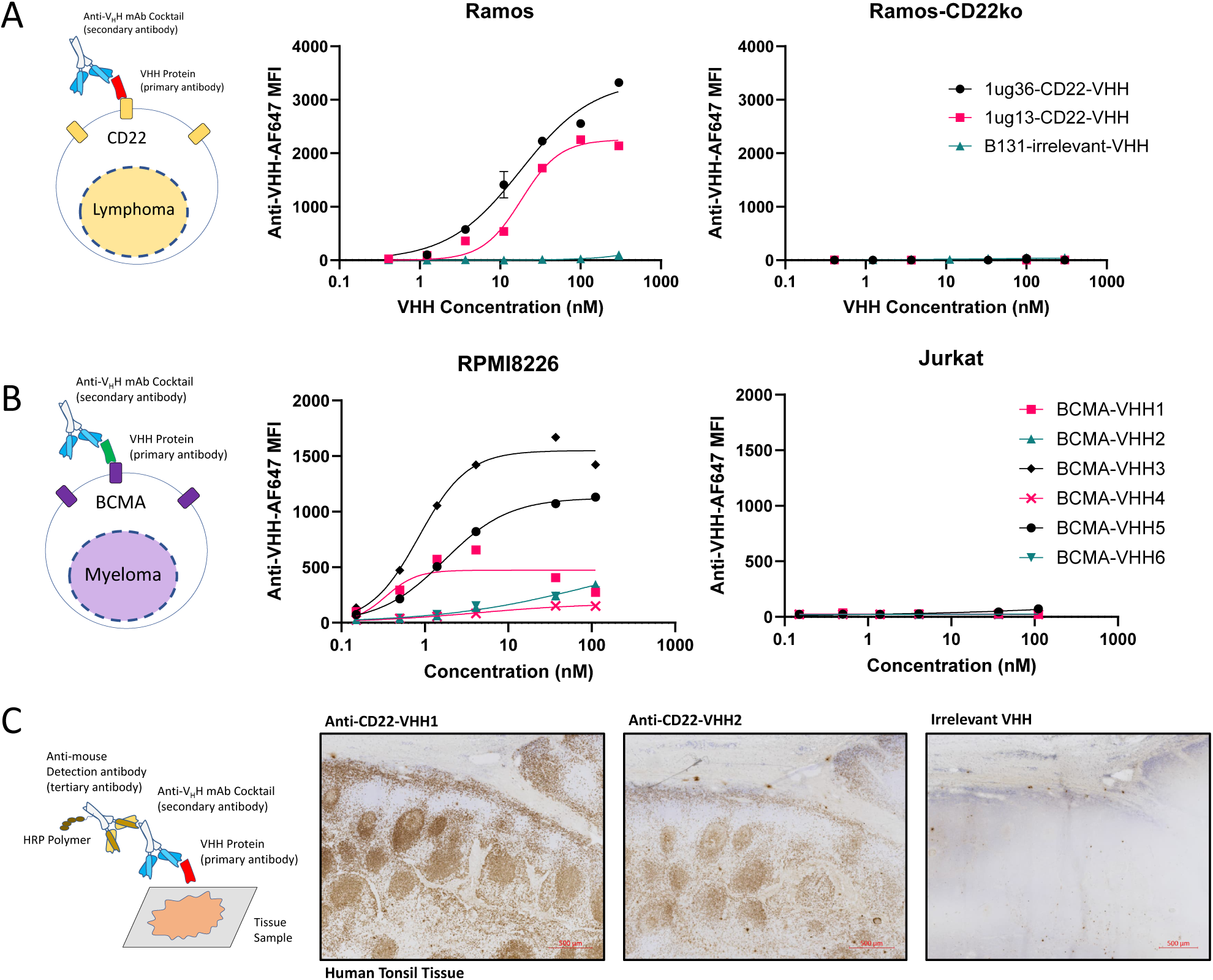
Anti-VHH mAbs can be used to assess cell and tissue binding of novel VHHs (A) As a demonstration of potential application for anti-VHH mAbs for assessment of VHH binding to cells, a human lymphoma cell line with high expression of CD22 (Ramos) or a similar line wherein CRISPR gene targeting was used to knockout CD22 expression (Ramos-CD22ko) were stained with purified CD22-specific VHHs (1ug36 or 1ug13 clones) or a control VHH of irrelevant specificity (B131). Cells were then washed and stained with a mix of AlexaFluor647 conjugated anti-VHH secondary mAb cocktail (3H12 and 2A3), and cells were examined via flow cytometry. Results demonstrate that VHH cell binding can be directly assessed using a secondary anti-VHH staining approach. (B) To demonstrate the general utility of this approach for VHHs of unknown sequence identity and cellular reactivity a panel of BCMA-specific VHHs was tested for reactivity to a BCMA+ multiple myeloma cell line (RPMI8226) or a BCMA-negative control cell line (Jurkat), followed by secondary staining with an anti-VHH cocktail. (C) To examine whether a similar approach can be used for assessment of tissue level reactivity of VHHs, human tonsillar tissue known to have a high density of CD22+ B cells was stained with CD22-specific VHH. Tissue were then stained with 3H12 anti-VHH mAb, and a tertiary anti-mouse HRP polymer detection reagent. Images show expected follicular tissue distribution consistent with CD22 specific reaction.

### Soluble Anti-VHH mAb Binding can Block VHH-CAR Reactivity

Having demonstrated that anti-VHH mAbs can bind to both soluble and CAR-linked VHH moieties, we next wanted to assess how soluble anti-VHH mAbs would affect VHH-CAR response to antigen expressing target cells. Jurkat cells with stable expression of a previously reported EGFR-specific VHH-CAR^11^ were co-incubated with EGFR-positive human H292 lung cancer cells (Figure 6A). EGFR-CAR Jurkat cells expressed high levels of the activation marker CD69 in the presence of H292 cells, whereas the addition of increasing doses of anti-VHH mAbs blocked CD69-upregulation with an IC_50_ of ∼0.1nM for the anti-VHH mAb mix (Figure 6B). Similarly, in primary CAR-T cells transduced with EGFR-VHH CAR lentivirus, soluble anti-VHH mAb treatment resulted in a dose-dependent blockade of CAR-T activation, as assessed by the CD69 and CD25 activation markers (Figure 6C). Furthermore, we observed potent inhibition of EGFR-VHH CAR-T cytolytic killing of EGFR-positive SKOV3 ovarian cancer cells with soluble anti-VHH mAb treatment (Figure 6D,E). These results show that anti-VHH mAbs can be employed to block antigen-specific VHH-CAR function.

**Figure 6:**
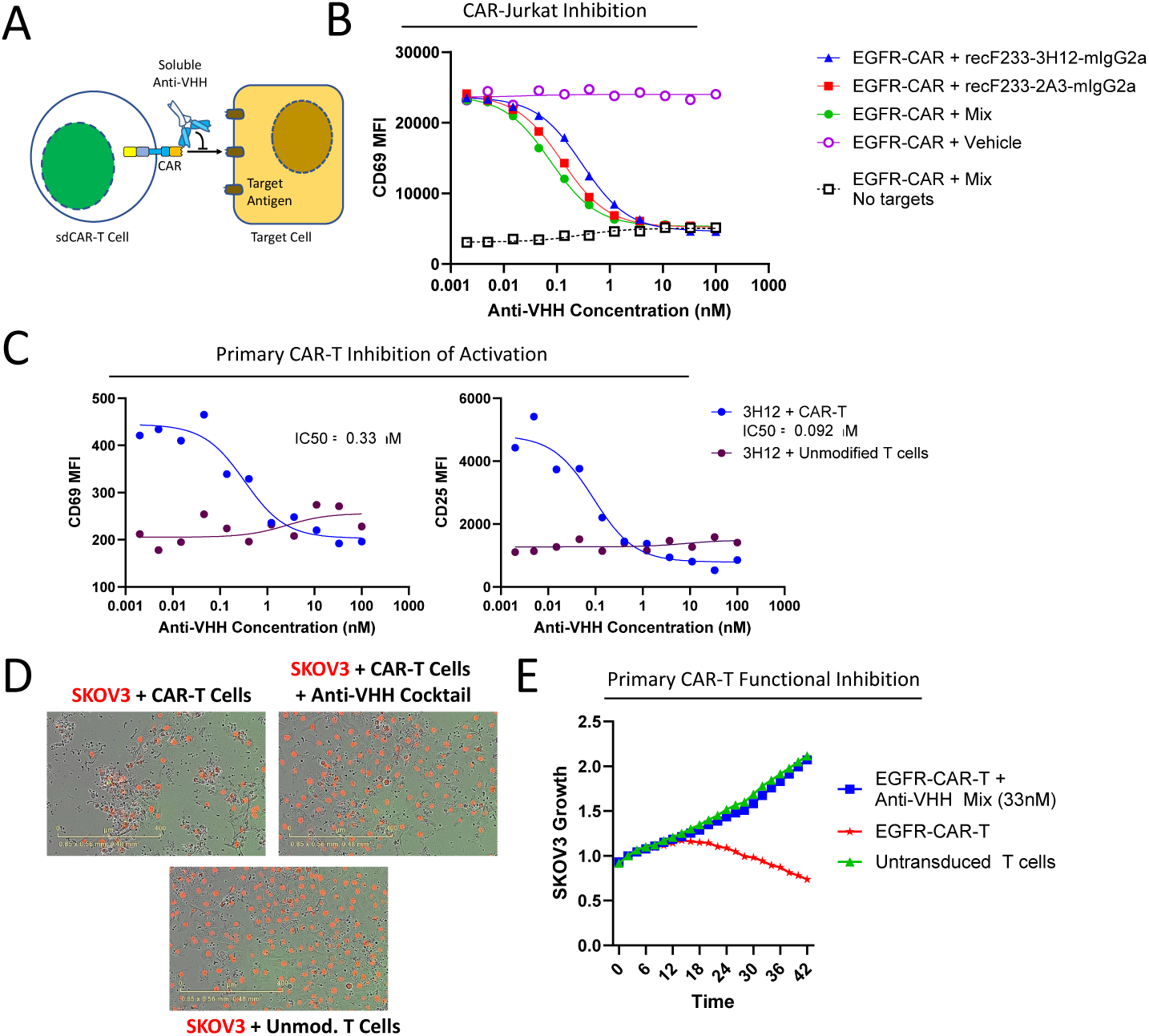
Soluble anti-VHH mAbs can block CAR-T cell responses (A-B) To test the effect of soluble anti-VHH mAbs, EGFR-specific VHH-CAR expressing Jurkat cells were combined with EGFR-expressing target cells and varying doses of anti-VHH mAbs separately or as a cocktail. CAR-Jurkat and target co-cultures with varying anti-VHH treatment dose were incubated for 48 hours before staining with anti-CD69 antibodies to assess effector cell activation via flow cytometry. Results show that CAR-Jurkat cells have a high level of activation with EGFR-expressing targets (open circles) but not without target cells (open squares). Anti-VHH mAbs alone or in combination resulted in a dose dependent blockade of CAR-Jurkat activation in response to EGFR-expressing target cells. (C) A similar experiment was performed with primary human CAR-T cells specific, wherein upregulation of both CD69 or CD25 activation markers were diminished in a dose-dependent manner with a soluble anti-VHH mAb. (D) Primary CAR-T cells in co-culture with red-fluorescent protein expressing EGFR-high SKOV3 target cells were examined for target cell survival via live fluorescence microscopy using an Incucyte device. Images taken after 42 hours in co-culture, show a clear loss of target cells when co-cultured with EGFR-specific VHH CAR-T cells (top left) but not unmodified human T cells (bottom). CAR-T mediated target cell killing was absent with soluble anti-VHH mAb treatment (right image). (E) Relative red fluorescent signal over time as measured via Incucyte shows that anti-VHH mAb treatment completely blocks CAR-T specific lysis of target cells.

### Immobilized Anti-VHH mAb Can Activate and Expand VHH-CAR Cells

We next wanted to assess whether immobilization through surface absorption of anti-VHH mAbs could be used to specifically expand CAR-T cells, similarly to what is commonly performed with CD3-specific mAbs. Upon exposure to various doses of plate-bound anti-VHH mAbs, we observed a dose-dependent upregulation of the T-cell activation marker CD69 in EGFR-VHH CAR-Jurkat cells but not in WT Jurkat cells (Figure 7A,B). Interestingly, anti-VHH mAb activation in this case resulted in lower magnitude of activation than with plate bound anti-CD3 mAb. To investigate the consistency of this activation across different VHH-CARs, we tested Jurkat cells with stable expression of 5 different CD22-specific VHH CARs, a murine anti-CD19-scFv CAR (FMC63) or a human CD22-scFv CAR (m971). While the anti-CD3 mAb OKT3 was able to activate all CAR-expressing cells and WT Jurkat cells as expected, plate-bound anti-VHH mAb resulted in varying activation of all VHH CARs but not scFv or WT Jurkat cells (Figure 7C). Generally, the 3H12 mAb resulted in a slightly more potent activation relative to 2A3, but a mixture of both mAbs resulted in activation of all Jurkat VHH-CAR cell lines, with slightly lower potency than plate-bound anti-CD3 OKT3 mAb.

**Figure 7:**
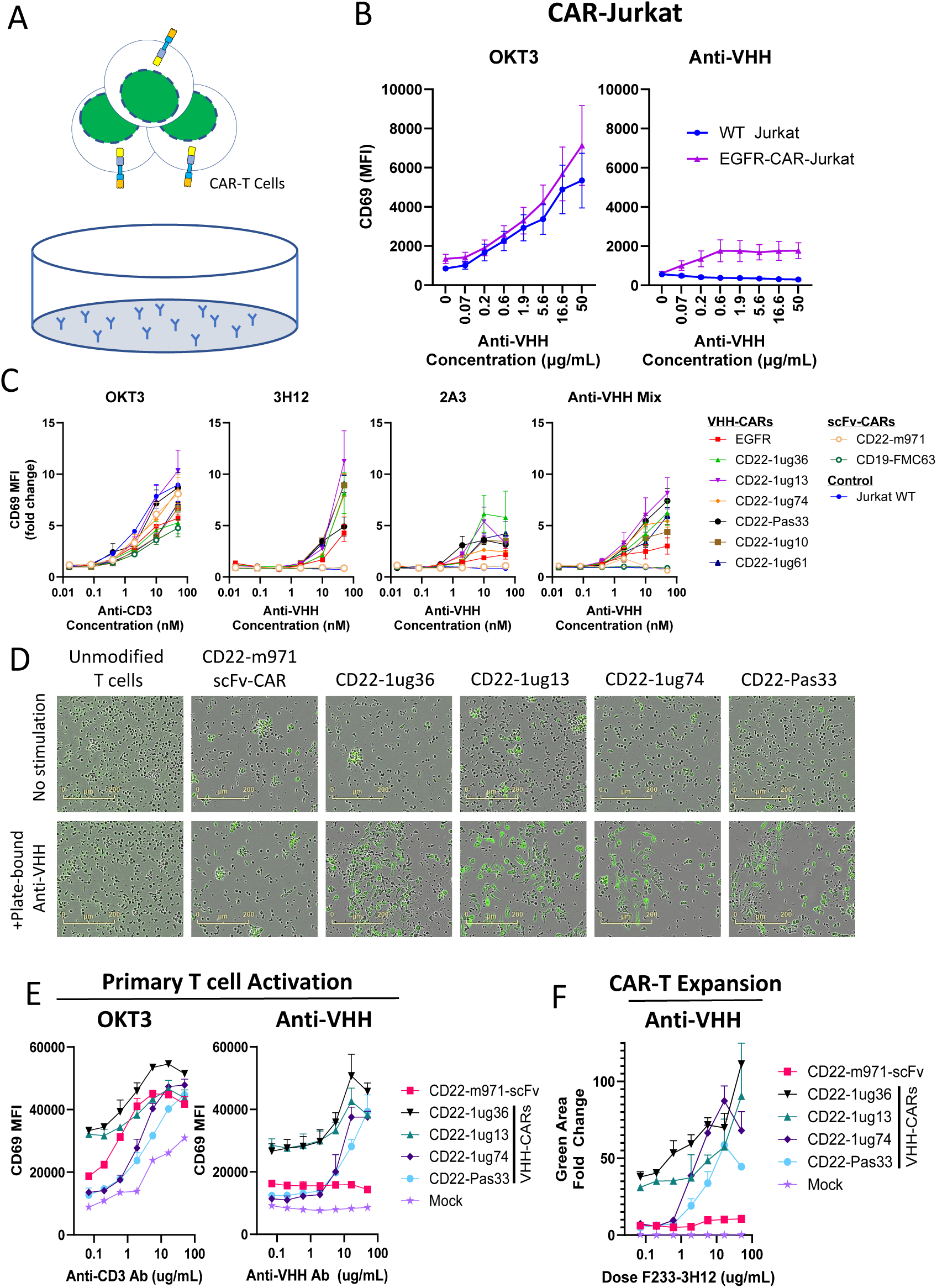
Immobilized anti-VHH mAbs can be used to activate and expand VHH-CAR-T cells (A) Schematic of experimental setup to test whether plate-bound anti-VHH mAbs were able to activate VHH-CAR expressing cells. (B) Varying concentrations of anti-VHH mAbs were absorbed to a 96-well plate overnight, before washing and addition of CAR-Jurkat cells for an additional 24 hours. Cells were then stained with CD69 antibody an assessed via flow cytometry. Whereas CD3-specific OKT3 antibody was able to strongly activate Jurkat cells regardless of CAR expression, only VHH-CAR expressing cells were activated by plate bound anti-VHH mAb. (C) To confirm this effect across varying cells expressing various VHH-CAR constructs, cells were stimulated with plate-bound 3H12, 2A3, a mix of both anti-VHH mAbs, or plate bound anti-CD3. Results show that all VHH-CAR expressing cells showed activation after incubation with plate-bound anti-VHH mAbs, whereas scFv-CAR expressing cells (FMC63- or m971-CARs) showed no activation. (D) A similar experimental setup was used to confirm that plate-bound anti-VHH mAbs can stimulate activation of primary human CAR-T cells. Images show a clear increase in size for human T cells expressing VHH CARs, but not for scFv-CAR-T cells. (E) Similarly VHH CAR-T cells show upregulation of the CD69 activation marker and (F) expansion of green fluorescent marker co-expressed with the CAR transgene. Overall, these results clearly demonstrate that plate immobilized anti-VHH mAbs can be used to activate and expand VHH-CAR expressing T cells.

To examine whether plate bound anti-VHH mAb could similarly activate healthy donor CAR-T cells, we generated CAR-T cells expressing 4 different CD22-VHH CARs or a control CD22-scFv CAR via lentiviral transduction. At day 12 after initial polyclonal activation using CD3/CD28 stimulation, cells were initially cryopreserved for later analysis. VHH-CAR or control cells were thawed and stimulated with plate-bound 3H12 mAb, resulting in a visible phenotypic change by 24 hours in VHH-CAR expressing CAR-T cells stimulated with anti-VHH mAb but not for scFv-CAR-T or untransduced control T cells (Figure 7D). Expression of the CD69 activation marker was increased in a dose-dependent manner for anti-VHH mAb stimulated VHH-CAR expressing cells but not scFv-CAR or control T cells (Figure 7E). Over time, anti-VHH mAb stimulated VHH-CAR-T but not control cells showed a strong increase in GFP signal, formed large clumps of cells, and began proliferating (Figure 7F, Supplemental Videos 1-6). At day 6 after anti-VHH mAb stimulation, cells were split to a low density in larger culture vessels with IL7 and IL15 cytokine supplementation. At day 22 anti-VHH mAb-expanded CAR-T products were assessed, revealing between 488 and 938-fold increase in total cell yield with >94% VHH-CAR expression. Anti-VHH mAb stimulation resulted in expansion of both CD8 and CD4 CAR-T cells, with a higher proportion of CD8 cells than the starting CAR-T product (Table 1). Overall, these data demonstrate that anti-VHH mAbs can be used to produce an expanded and purified VHH-CAR expressing cell product.

**Table 1:** Expansion of VHH CAR-T samples using plate-bound Anti-VHH mAb.

|  | At CAR-T harvest |  |  | After Anti-VHH mAb Expansion |  |  |  |  |
| --- | --- | --- | --- | --- | --- | --- | --- | --- |
|  | %CAR+<br>(GFP) | %CD8+ | %CD4+ | %CAR+<br>(GFP) | %CD8+ | %CD4+ | Total Cells | Fold Growth |
| Unmodified T cells | 0.38 | 22.1 | 60.5 | 0.02 | 4.7 | 19.1 | 315 000 | 20 |
| CD22-m971 scFv | 32.0 | 11.2 | 58.2 | 56.5 | 42.3 | 5.27 | 680 000 | 43 |
| CD22-1ug36 | 24.4 | 7.8 | 66.9 | 97.2 | 26.7 | 51.9 | 12 800 000 | 800 |
| CD22-1ug13 | 33.6 | 8.6 | 62.8 | 96.0 | 28.6 | 47.6 | 15 000 000 | 938 |
| CD22-1ug74 | 36.0 | 12.7 | 78.5 | 94.8 | 37.4 | 23.9 | 7 800 000 | 488 |
| CD22-Pas33 | 36.0 | 12.3 | 60.5 | 94.7 | 14.8 | 52.3 | 11 100 000 | 694 |

### Anti-VHH mAb Expanded VHH-CAR-T Cells Maintain Antigen-Specific Functionality and Long-term Response Potential

We finally tested the functionality of CAR-T cells after anti-VHH mAb expansion using co-culture with varying target cells (Figure 8A). First, to examine the short-term response of CAR-T cells with or without anti-VHH mAb activation, CAR-T cells were placed in co-culture with CD22+ Ramos cells at varying effector to target ratios for 7 days. We observed that anti-VHH mAb expanded cells maintain similar or better killing of CD22+ target cells as non-activated CAR-T cells (Figure 8B, Supplemental Figure 3A). To test the persistence of cells expanded with anti-VHH mAbs, we examined growth in longer-term exhaustion-inducing conditions, wherein additional target cells were added to co-cultures 2 times per week, and cultures were split weekly for 3 weeks. As expected, we observed that VHH-CAR-T cells display exhaustion under these conditions, with red-fluorescent target cell dominating the co-cultures at weeks 2 and 3 (Figure 8C, Supplemental Figure 3B). Interestingly, a benchmark m971 scFv-based CAR-T which has been engineered for increased tonic signaling^17^ shows only transient exhaustion, regaining capacity to restrict target cell growth at week 3 of co-culture. Examining the long-term response in anti-VHH mAb expanded VHH-CAR-T cells, we were surprised to also observe resistance to exhaustion, with CAR-T dominated co-cultures at week 3 with anti-VHH mAb-activated effector cells (Figure 8C, Supplemental Figure 3B). Importantly, anti-VHH mAb-expanded CAR-T cells do maintain antigenic selectivity, as no long-term restriction of CD22-knockout lymphoma or CD22-negative lung cancer cells was observed. In conclusion, we report that anti-VHH mAb expansion produces CAR-T cells that maintain strong short-term functionality and have potential for long-term antigen-specific responses.

**Figure 8:**
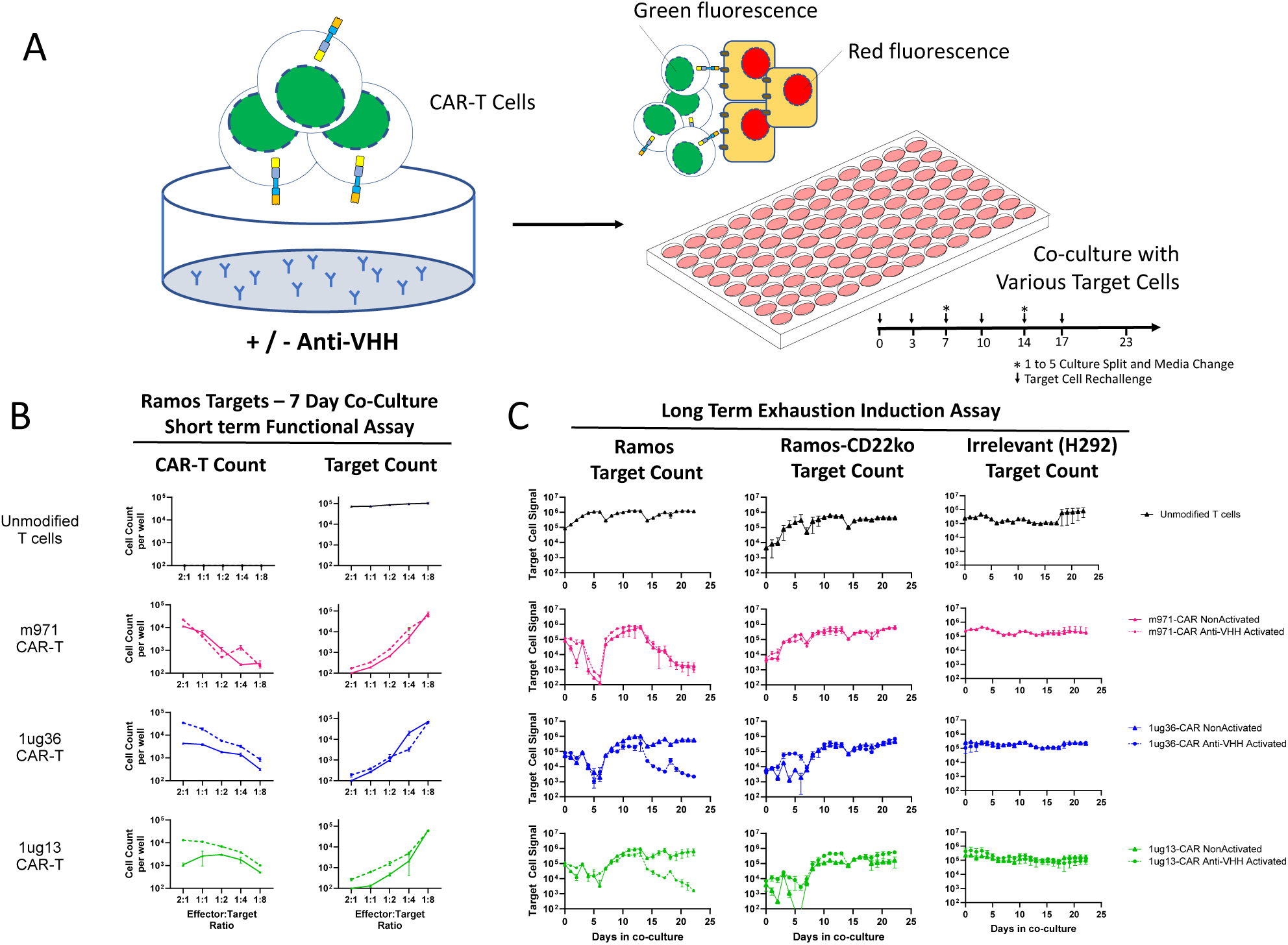
Functional testing of anti-VHH Activated CAR-T Cells (A) Schematic of experimental setup to activate VHH CAR-T cells with plate-bound anti-VHH mAbs, followed by functional assay using co-culture with various target cells. (B) The short-term response to CD22+ Ramos target cells was examined across a range of effector to target ratios, wherein the number of red-fluorescent target cells and green-fluorescent CAR-T cells were enumerated using flow cytometry after 7 days of co-culture. (C) Co-cultures of 20 000 CAR-T cells with an equal number of CD22+ Ramos, CD22-knockout Ramos, or CD22-negative H292 lung cancer target cells. To drive CAR-T exhaustion, additional target cells were added to cultures at days 3, 7, 10, 14, and 17; 1 to 5 media exchange and cell culture split were also performed on days 7 and 14. During this entire period, growth of red-fluorescent target cells were monitored via live regular fluorescent microscopy. All graphs show the response of non-activated CAR-T cells (solid lines) versus anti-VHH activated cells (broken lines) from a single experiment.

## Discussion

In this work, we report the development of a standardized Ab reagent for directly labelling and manipulating VHH CAR-T cells. In recent years, we have reported on our pre-clinical development of novel camelid VHH-based CARs targeting CD22^10^ and EGFR^11^, and we are actively working on similar VHH-CAR therapeutics targeting BCMA, CD20, mesothelin, and other cancer-associated antigens. The broad reactivity of the anti-VHH mAb labelling reagents allows us to apply this tool across diverse VHH-CAR discovery projects, directly confirming surface expression during early VHH-CAR high-throughput screening^18^, and obviating the need for a fluorescent marker in CAR-screening plasmids.

The anti-VHH mAbs we have developed are functionally analogous to anti-idiotypic mAbs against FMC63, the most common CD19-scFv used in CARs^19^, aptamers specific for the FMC63 element of CD19-CARs^20^, or fluorophore-tagged antigens. In contrast to such Ab or antigen specific strategies, the anti-VHH mAbs reported here can be applied for detecting expression of VHH CAR targeting any antigen. Other more generalized strategies for labeling CAR molecules such as using fluorescent protein A, anti-linker Abs, and polyclonal anti-variable domain Abs^21^, are less applicable to VHH-CARs; protein A reacts with only a subset of VHH molecules and VHH-CARs lack the typical G4S or Whitlow linkers used in scFv-based CARs. While polyclonal anti-VHH Abs are now available, the lack of consistency and characterization of defined products can make pre-clinical and clinical development using such reagents challenging. The data presented here demonstrates that our dual-clonal anti-VHH mAb cocktail (3H12 and 2A3 mix) is a stable and consistent product even after fluorophore conjugation which could be applied in various high-quality release assays of pre-clinical and clinical grade VHH-CAR cell and viral products.

In order to maximize the potential utility of our anti-VHH mAbs, we performed extensive characterization. Our epitope mapping results delineate two unique CDR-distal FR epitopes recognized by these mAbs which imparts advantages for the broader application of the mAb cocktail. Deuterium exchange results along with competitive SPR inhibition for mAb 3H12 and 2A3 support distinct binding epitopes that are not entirely mutually exclusive and suggests close proximity to one another. The targeted epitopes here are contrary to a previously reported CDR-distal anti-VHH Fab domain^22^ wherein binding involved a C-terminal tag. Amino acid sequence alignment of non-reactive and reactive VHHs revealed that there were several mutations unique to those VHHs not recognized by the mAb 3H12 including a serine residue at position 11 (IMGT numbering) in the VHH FR1 domain, wherein mutational reversion (S11L) fully restored 3H12 binding. While we were not able to pinpoint key interaction residues for the 2A3 mAb, our results clearly demonstrate that a dual-clonal anti-VHH mAb mix is applicable for VHH binding to the majority of VHHs, with the mixture of mAbs recognizing 75 out of 76 llama-derived VHHs tested. Based on our overall assessments of anti-VHH mAb epitopes, we expect that the cocktail would also likely be cross-species reactive for other VHHs, such as those derived from camel and alpaca, due to the conserved FR regions across species^23^; however further work will be necessary to confirm this.

In addition to surface labeling, we found that anti-VHH mAbs can block VHH-CAR reaction against antigen positive target cells. A similar effect has been previously reported with anti-idiotypic Abs against CD19-CAR^19^, and is analogous to the effect of anti-CD3 mAb blockade of T-cell receptor mediated immune responses^24^. We believe this represents the first Ab product which has been demonstrated to be useful for direct blockade of VHH-CAR activation. Given that anti-VHH mAb epitopes are CDR-distal, and reactivity does not disrupt VHH-antigen interaction in cell labeling experiments, we infer that the mechanism of VHH-CAR disruption is likely through inhibition of CAR clustering or other downstream signaling mechanisms. These CAR-blocking mAbs could prove useful for investigating the nature of VHH-CAR signaling, or to identify the optimal signaling magnitude and timing to increase CAR therapeutic efficacy *in vitro* or *in vivo*. Generating an infusible product derived from these anti-VHH mAbs could also have clinical utility, as it could conceivably allow for rapid and titratable blockade of VHH-CAR activity if and when toxicity is observed.

Another intriguing application of the anti-VHH mAbs reported here is the specific expansion of VHH-CAR expressing T cells. Various techniques for expansion of CD19 CAR-T cells have been established previously, including stimulation with antigen-expressing artificial antigen presenting cells^25,26^, antigen-bearing vesicles^27,28^, and multimerized target antigen^29^. While these strategies are proven to generate highly functional CAR-T products, they are target-antigen specific and thus could not be broadly applied to novel CAR-T products. In contrast, stimulation with plate-bound anti-VHH mAbs is a target-agnostic approach capable of producing >500-fold CAR-T expansion, while maintaining CD4/CD8 heterogeneity and antigen-specific CAR-T functionality.

It is well documented that repeated stimulation through CD3 can lead to functional exhaustion of T cells (reviewed in ^30^), and recent developments favour shortened manufacturing strategies with less CAR-T expansion to improve long-term persistence of CAR-T therapies^31,32^. Given this, we were concerned whether anti-VHH mAb expansion could lead to functional exhaustion of VHH-CAR-T cells. In contrast to our initial hypothesis, our *in vitro* assessments indicate that anti-VHH mAb expanded cells maintain strong target specific activity and appear to be resistant to exhaustion inducing conditions. While these initial results are encouraging with regard to the use of anti-VHH mAbs to produce larger numbers of functional CAR-T cells, further experiments will be needed to characterize cellular signaling induced through stimulation with immobilized anti-VHH mAbs and the downstream transcriptional effect in CAR-T cells. In pursuit of this, we are currently developing a more refined and consistent method to expand VHH-CAR-T cells using bead-bound anti-VHH mAbs and investigating the potency of expanded CAR-T products in xenograft mouse models.

Overall, we have demonstrated that the new anti-VHH mAbs presented here are highly useful tools for labeling and manipulating VHH-CAR expressing cells, which could be of significant utility to the growing body of researchers working towards the development of more advanced VHH-CAR therapeutics.

## Materials and Methods

### Mouse Immunization

Four six-week-old female A/J mice (The Jackson Laboratory, USA, Cat#000646) were bled (pre-immune serum) and injected intraperitoneally and subcutaneously with 100 µg of VHH antigen emulsified in TiterMax adjuvant (Cedarlane Labs, Canada, Cat#R-5) at day 0 and in PBS without adjuvant at day 26. Blood was collected in microvette CB 300Z (Sarstedt, Canada, Cat#16-440) at day 33, and serum was stored at −20°C until further use. Pre- and post-immune sera titer of animals immunized with VHH antigen were assessed by ELISA. Unless otherwise stated, all incubations were performed at room temperature. Briefly, half-area 96-well plates (Costar, USA, Cat#3690) were coated with 25 µl per well of VHH at 5 µg/ml in PBS and incubated overnight at 4°C. Microplates were washed three times in PBS and blocked for 30 minutes with PBS containing 1% bovine serum albumin (BSA, Sigma, USA, Cat#A7030). Blocking buffer was removed and 25 µl of serial dilutions of sera samples were added. After a 2 hour incubation, microplates were washed 4 times with PBS-Tween 20 0.05% and 25 µl of a 1/5000 dilution of alkaline phosphatase conjugated goat anti-mouse IgG (H+L) (Jackson ImmunoResearch, Cat#115-056-062) in blocking buffer was added. After a 1 hour incubation, microplates were washed 4 times and 25 µl of p-nitrophenyl phosphate (pNPP) substrate (Sigma, USA, Cat#N7653) at 1 mg/ml in carbonate buffer at pH 9.6 was added and further incubated for 30 minutes. Absorbance was read at 405 nm using a SpectraMax plate reader (Molecular Devices, USA). All pre-immune bleeds were negative and all post-immune bleeds were very strong (above 1/12800) on immunogen.

### Hybridoma Generation

After 2-3 months, an i.p. booster injection (100 µg of FC5-MOD protein produced in house, in PBS) was done 3 days prior to fusion experiment. Spleen cells were harvested in Iscove’s Modified Dulbecco’s medium (IMDM, Gibco, USA, Cat#31980-030) and fused to NS0 myeloma cell line using polyethylene glycol. Spleen cells and myeloma cells were washed in IMDM, counted in RBC lysing buffer (Sigma, USA, Cat#7757-100ML) and mixed together at a 5:1 ratio. Pelleted cells were fused together by adding 1 ml of a 50% solution of PEG 4000 (EMD-Millipore Cat#9727-2) in PBS preheated at 37°C drop-wise over 1 minute, and incubated at 37°C for an additional 90 seconds. The reaction was stopped by addition of 30 ml of IMDM at 22°C over a period of 2 minutes. After a 10 minute incubation, freshly fused cells were spun at 233g for 10 minutes. Cells were washed once in IMDM supplemented with 10% heat inactivated FBS (Sigma Cat #F1051).

Following fusion, cells were suspended at a concentration of 2×10^5^ input myeloma cells per ml in HAT selection medium (IMDM containing 20% heat inactivated FBS, penicillin-streptomycin (Sigma, USA, Cat#P7539), 1 ng/ml mouse IL-6 (Biolegend, USA, Cat#575706), HAT media supplement (Sigma, USA, Cat#H0262) and L-glutamine (Hy-Clone Cat#SH30034.01) and incubated at 37°C, 5% CO2. The next day, hybridoma cells were washed and suspended at a concentration of 2-3 × 10^5^ input myeloma cells per ml in semi-solid medium D (StemCell Technologies, Canada, Cat. No. 03804) supplemented with 5% heat inactivated FBS, 1 ng/ml mouse IL-6 and 10 µg/ml FITC-F(ab’)2 Goat anti-mouse IgG (Jackson Immunoresearch, USA, # 115-096-071). The cell mixture was plated in Omnitray dish (Nunc, USA, Cat#242811) and further incubated for 6-7 days at 37°C, 5% CO2. Fluorescent secretor clones were then transferred using a mammalian cell clone picker (ClonepixFL, Molecular Devices, USA) into sterile 96-well plates (Costar, USA, #3595) containing 200 µl of IMDM supplemented with 20% heat inactivated FBS, penicillin-streptomycin, 1 ng/ml mouse IL-6, HT media supplement (Sigma, USA, Cat# H0137) and L-glutamine and incubated for 2-3 days at 37°C, 5% CO2.

### ELISA on Hybridoma Supernatants

Hybridoma supernatants were screened by enzyme linked immunosorbent assay (ELISA) to detect specific binders. To this end, 96-well half-area plates (Costar, USA #3690) were coated with 25 µl of FC5-MOD or FC5 derivatives FC5-VHH or FC5-hFc-1X0, or P257 (unrelated VHH) or EG2-hFc-X2 (unrelated VHH) or BSA (negative control) at 5 µg/ml in PBS and incubated overnight at 4°C. Microplates were washed 3 times with PBS, blocked with PBS-BSA 1%, and 25 µl of hybridoma supernatants were added and incubated at 37°C, 5% CO_2_ for 2 hours. Plates were washed 4 times with PBS-Tween 20 0.05% and incubated for one hour at 37°C, 5% CO_2_ with 25 µl of secondary antibody alkaline phosphatase conjugated F(ab’)2 goat anti-mouse IgG Fc-gamma specific (Jackson Immunoresearch, USA, # 115-056-071) diluted 1/3000 in blocking buffer. After 4 washes with PBS-Tween 20 0.05%, 25 µl of a 1 mg/ml pNPP substrate solution was added and further incubated for one hour at 37°C. OD405nm measurements were done using a microplate reader (Spectramax® 340 PC, Molecular Devices, USA). Hybridoma supernatants were further analyzed for positive reactivity to various VHHs: VHH immunogen, 1ug13, and 1ug36; and for negative reactivity to human antibodies using protX-hIgG (negative control).

### Recombinant Antibody Production and Purification

The VH and VL of each mAbs were sequenced by Sanger sequencing, determining the CDRs sequence were determined using Kabat CDR numbering system. The VH and VL regions from 2A3 and 3H12 were cloned as fusions with mouse IgG2a/kappa constant regions (mouse IgG2a heavy chain and mouse kappa light chain, respectively) into the in house pTT109 plasmid vector. Chinese hamster ovary (CHO) cells were then transfected with VL and VH containing constructs (1:1 ratio). Conditioned medium (CM) was harvested on day 7, recombinant antibodies were purified on Protein A (HiTrap MabSelect SuRe, Cytiva, USA) connected to an AKTApure or an AKTAavant, buffer exchanged in DPBS on HiPrep™ desalting column (Cytiva, USA) connected to an AKTAexplorer, sterile-filtered, quantitated using a Nanodrop 2000 system, and evaluated by UPLC-SEC and SDS-PAGE. AF647 labeling was performed using a commercial kit (Invitrogen, USA, Cat#A20173), according to manufacturer’s instructions.

### VHH Expression and Purification

VHHs were expressed from pMRO.BAP.his plasmid in *E. coli* BL21(nG3) using standard conditions. VHHs were isolated from the bacteria cell pellet by sonication in IMAC binding buffer, and lysates were cleared by centrifugation and filtration through 0.2 mm PES filter. Each VHH was affinity-purified using HisTrapHP columns (Cytiva, USA) and gradient imidazole elution on an AKTAxpress (GE Healthcare, USA). His-purified VHHs were buffer exchanged to PBS using centrifugal filters (Amicon, USA) and further purified using a Superdex 75 Increase 10/300 GL size exclusion chromatography (SEC – Cytiva, USA) column connected to an AKTApurifier (GE Healthcare, USA).

### ELISA Detection of VHHs with Anti-VHH mAbs

VHHs were diluted to 10 µg/mL in PBS and 1 µg VHH was absorbed to each uncoated ELISA well at 4°C overnight. All subsequent steps were performed at room temperature. Wells were blocked with 5% (w/v) skim milk in PBS for 1 hour; followed by incubation with 1:5000 anti-VHH mAb (3H12 and 2A3) in blocking buffer for 1 hour; 3× washing with PBS + 0.05% Tween 20; incubation with 1:5000 HRP conjugated donkey anti-mouse-IgG (Jackson Labs, Cat#715-035-150) in blocking buffer for 1 hour; 3× washing with PBS + 0.05% Tween20; and incubation with TMB substrate (Abcam, USA) for 5 minutes before stopping the reaction with 1 M phosphoric acid.

### Surface Plasmon Resonance Binding Assays

All SPR experiments were conducted at 25°C on a Biacore T200 instrument (Cytiva, Canada) in PBST running buffer (135 mM NaCl, 2.7 mM KCl, 4.3 mM Na_2_HPO_4_, 1.4 mM KH_2_PO_4_, 3 mM EDTA, 0.05% (v/v) Tween 20, pH 7.2). For measurement of VHH monovalent affinities to 3H12 and 2A3 mAbs, approximately 2000 response units (RUs) of goat-anti-mouse IgG Fc (Jackson ImmunoResearch, USA, Cat#115-005-071) were amine coupled on a CM5 series S sensor chip (Cytiva) in 10 mM acetate buffer, pH 4.5, prior to capture of 3H12 or 2A3 mAbs (100 µg/mL) at 10 µL/minute flow rate and 60 s contact time. VHHs, previously purified by SEC, were injected at concentrations ranging from 2 – 1000 nM over captured 3H12 and 2A3 surfaces at a flow rate of 20 µL/minute, with 180 s of contact time and allowed to dissociate for 600 s. Surfaces were regenerated with 10 mM glycine, pH 1.5, at 30 µL/minute for 120 s. Reference flow cell subtracted single-cycle kinetics data was fit to a 1:1 interaction model using BIAevaluation 3.2 software (Cytiva, USA). For SPR competition experiments, a VHH that was recognized by both 3H12 and 2A3 mAbs was amine coupled (130 RUs) on a CM5 series S sensor chip as above. Next, 3H12 (100 nM) was injected at 20 µL/minute for 120 s followed immediately by injection of a mixture of 3H12 (100 nM) + 2A3 (100 nM) for 120 s. The reverse order was also performed (2A3 at 100 nM, followed by a mixture of 2A3 + 3H12, both at 100 nM). Surfaces were regenerated with 1 M MgCl_2_ at 10 µL/minute for 10 s.

### VHH Yeast Surface Display and anti-VHH mAbs binding

Recombinant anti-VHH mAbs were examined for reactivity with truncated mutants of CD22-specific VHH 1ug36 generated as yeast-surface displayed fusion proteins (Aga2-HA-(VHH)-MYC). Yeast cells were then grown under the inducing condition in galactose media. The binding of the mAbs to yeast cells was performed using a whole yeast cell ELISA. Briefly, induced yeast cells were washed with 1× PBS, and 1 × 10^6^ cells were transferred per well to 96-well filter plate (#MSDVN6550 MultiScreen DV Filter Plate, Millipore). After adhering yeast cells, PBS was removed by vacuum aspiration and 200 µl of blocking solution (PBS with 2 % BSA and 0.05 % Tween 20, Sigma Aldrich, USA) was added, and plates were incubated for 45 minutes at room temperature with shaking. Blocking solution was removed by vacuum aspiration and 100 µl of biotinylated anti-VHH mAbs (100-250 nM), or irrelevant mouse monoclonal anti-c-myc antibody (9E10) at 3 µg/ml (Sigma) was added and incubated for 1 hour at room temperature with shaking. Cells were washed 4× with 200 µl per well PBS + 0.05 % Tween 20. Detection of biotinylated VHHs was performed using 100 µl of HRP-conjugated streptavidin (Jackson # 016-030-084) at 1/10 000 dilution in 1 % BSA + 1× PBS + 0.025% Tween 20, and MYC-tag was detected with 100 µl per well of HRP-F(ab)2 fragment goat anti-mouse IgG (H+L) (Jackson ImmunoResearch, Cat #115-036-062) at 1/ 5000 in the same binding solution. Mixture was incubated for 1 hour at room temperature with shaking, then washed 4 × with PBS + 0.05 % Tween 20, followed by adding 100 µl per well of TMB substrate solution (Thermo Fisher, USA, Cat#34021) and incubated for 10 minutes at room temperature, before stopping with 0.2 M sulfuric acid. Absorbance at 450 nm (OD450) was measured using a Tecan Infinite M1000 Pro plate reader.

### HDX Mapping

For HDX-MS, purified sdAb-1ug36 protein was equilibrated with a saturating concentration of each anti-VHH mAb (1:1 molar ratio). Deuterium labelling of the mAb-VHH complexes or VHH (1ug36 clone) alone was initiated by a two-fold dilution with deuterated buffer (PBS in 90% D_2_O). A controlled time period of either 0.5 minute or 3 minutes was allowed before the reaction was quenched by a 5-fold dilution with 1 M guanidine HCl 40 mM TCEP in 0.1% formic acid at 4 °C. Samples were then trapped (ACQUITY UPLC BEH C18 VanGuard Pre-column, 130Å, 1.7 µm, 2.1 mm X 5 mm), washed, and digested with immobilized Nepenthesin-II (Affipro) at 100 µL/minute for 5 minutes (2 °C). Peptides were eluted from an Acquity UPLC column (PST BEH C18, 130A, 1.7um, 100umx100mm) with a 1-35% acetonitrile gradient (0.23% formic acid) at 40 µL/min. Data was collected on a Synapt G2-Si mass spectrometer with ion mobility enabled (Waters). A peptide map was generated with a database search (Mascot) of a data-dependent acquisition of an unlabelled 1ug36 VHH sample. Deuteration was assigned in triplicate for 41 peptides covering 81% of the sdAb-1ug36 sequence using HDExaminer and Mass Spec^33^ (Supplemental Figure 2, Supplemental Table 2). Differential HDX was assessed by subtracting the deuterium content of the unbound (1ug36) from the mAb-VHH complex (2A3 or 3H12 + 1ug36). Significance was assessed using a two-state Welch’s T-test with a 1-p threshold of 0.98 and a 3x standard deviation cut-off. Differences (ΔD = D_mAb+VHH_ – D_VHH_) in absolute (Δ #D) and relative (Δ % D) deuterium content are shown for each 1ug36 peptide. Relative data is normalized to the theoretical maximum deuterium of each peptide.

### Cell Binding and Immunohistochemistry (IHC)

Highly pure CD22- or BCMA-specific VHHs, and a control VHH specific for an irrelevant bacterial toxin (B131; as reported in ^34^), were produced using a standard bacterial expression system and metal affinity column purification. Myeloma (RPMI8226) or lymphoma (Ramos), or a modified cell line wherein CRISPR technology has been employed to specifically disrupt CD22 expression, were combined with varying concentrations of these purified VHH proteins. Cells were then washed and stained with the mixed AF-647 labelled 2A3 and 3H12 mAbs as a secondary antibody. Binding was then assessed via flow cytometry using a LSRFortessa flow cytometer (BD Bioscience, USA). Immunohistochemistry was performed using human tonsillar tissue which was fixed using methanol/Acetone. Tissue sections were then stained with primary CD22 VHH1 or VHH2, washed, and probed with a secondary anti-VHH mAb. This was then washed again, and stained with a tertiary anti-mouse-HRP polymer reagent as per manufacturer recommendation (Leica Biosystems, Canada, Cat#DS9800).

### CAR-Jurkat Activation or Inhibition Assays

For anti-VHH mAb inhibition assay, EGFR-CAR Jurkat cells were generated using lentiviral transduction as previously reported^11^. EGFR CAR-Jurkat were then placed at 37°C for 48 hours in co-culture with EGFR-positive SKOV3 human lung cancer cells, in the presence of decreasing concentrations of the anti-VHH mAbs 2A3 and 3H12, alone or in combination, or vehicle control. For activation studies, varying doses of anti-VHH mAbs were absorbed onto 96-well flat bottom cell culture plates (Corning, USA, Cat#3595) overnight in PBS. Jurkat cells with stable expression of various CD22 VHH-CAR or scFv-CARs as previously reported^10^ were then placed on the wells and incubated at 37°C and 5% CO_2_ overnight. Cells were stained with anti-CD69-APC (BD Bioscience, USA, Cat#555533), washed, and examined via flow cytometry.

### CAR-T Generation, Anti-VHH mAb Inhibition or Expansion, and Functional Testing

EGFR or CD22 primary CAR-T cells were generated from healthy PBMC or donor apheresis product respectively as previously described^11,15^. For inhibition studies, EGFR CAR-T cells were placed at an E:T ratio of 1:1 with red fluorescent protein expressing (Nuclight Lenti-Red, Sartorius, USA) SKOV3 cells in the presence of absence of various doses of anti-VHH mAbs. The relative growth of CAR-T and target cells was then monitored over 42 hours using an Incucyte device (Sartorius, USA). For anti-VHH mAb expansion studies, 20 000 CD22 CAR-T cells were placed in a 96-well flat bottom plate which was coated with varying doses of anti-VHH mAbs. Cells were observed regularly using live fluorescence microscopy via Incucyte and flow cytometry at day 7. After 6 days in culture, anti-VHH mAb activated CAR-T cells were also transferred to larger growth vessels to allow for full proliferation.

For the functional experiments testing with or without plate-bound anti-VHH mAb stimulation, cryopreserved CD22 CAR-T cells generated from healthy donor apheresis product as previously reported were used^15^. CD22 CAR-T cells or matched untransduced control T cells were thawed and allowed to recover for 24 hours before stimulating with plate-bound anti-VHH mAbs in a 24-well plate as described in the text. Cells were incubated in Immunocult XF media supplemented with IL-7 and IL-15, in 24-well tissue culture plates coated overnight with or without anti-VHH mAbs. Cells were allowed to expand for 12 days, with removal of excess cells to tissue culture flasks with no anti-VHH mAb as cells reached confluency, maintaining between 2.5 to 5 × 10^5^ cells per mL throughout the expansion period. For functional testing, cells were plated with target cells stably expressing Nuclight-Lenti-Red (Sartorius USA), with varying target cell types, and at varying effector to target ratios as described in the text. The relative growth of target and effector cells was examined using live fluorescence microscopy (Incucyte S3 device, Sartorius, USA). Every 2 to 3 days, additional target cells were added at equivalent numbers to initial challenge. Every 7 days, cultures were split 1 to 5 with fresh media simultaneously with additional target cells. At weekly splits, flow cytometric analysis was used to enumerate and characterize co-cultures.

---

**Supplemental Table 1:** ELISA results for screening of mouse anti-VHH hybridoma supernatants for VHH-reactivity and human-IgG reactivity. Hybridoma supernatants generated as described in Figure 1 were tested against surface absorbed VHHs (column 3-5) or total human IgG protein (column 6) for reactivity. Hybridoma supernatants were reacted with test antigens followed by washing and detecting with anti-mouse secondary antibodies. Results were used to select anti-VHH mAb candidates screened for reactivity to VHH-CAR cells as described in figure 1.

|  | mAb clone ID | Coating antigen (ELISA) |  |  |  |
| --- | --- | --- | --- | --- | --- |
|  |  | VHH 1 | VHH2 | VHH3 | hlgG1 antibody |
| 1 | F233-2A3-1 | 1.857 | 1.450 | 2.290 | 0.014 |
| 2 | F233-2D2-1 | 1.660 | 1.776 | 2.018 | 0.038 |
| 3 | F233-2D5-1 | 1.420 | 1.329 | 1.614 | 0.012 |
| 4 | F233-2E10-1 | 1.735 | 1.683 | 2.260 | 0.028 |
| 5 | F233-2G2-1 | 1.337 | 1.293 | 1.617 | 0.011 |
| 6 | F233-3F12-1 | 1.900 | 2.024 | 2.595 | 0.042 |
| 7 | F233-3G11-1 | 1.730 | 2.080 | 2.359 | 0.040 |
| 8 | F233-3H12-1 | 1.736 | 1.762 | 2.314 | 0.012 |
| 9 | F236-2A4-1 | 0.147 | 1.789 | 1.339 | 0.022 |
| 10 | F236-3C7-1 | 0.090 | 2.199 | 2.263 | 0.024 |
| 11 | F236-3A8-1 | 1.680 | 1.668 | 1.860 | 0.021 |
| 12 | F236-3E5-1 | 0.150 | 1.935 | 2.020 | 0.016 |
| 13 | F236-4B2-1 | 1.072 | 1.916 | 1.935 | 0.023 |
| 14 | F236-4E1-1 | 0.191 | 1.873 | 1.947 | 0.023 |
| 15 | F258-1B2-1 | 1.978 | 0.139 | 0.025 | 0.009 |
| 16 | F258-1C5-1 | 2.202 | 0.919 | 0.486 | 0.190 |
| 17 | F258-1H5-1 | 1.129 | 0.045 | 1.816 | 0.005 |
| 18 | F258-2A11-1 | 1.398 | 1.090 | 0.032 | 0.012 |
| 19 | F258-2B10-1 | 1.108 | 0.021 | 1.752 | 0.005 |
| 20 | F258-2C9-1 | 1.543 | 1.258 | 0.008 | 0.004 |
| 21 | F258-2E8-1 | 0.037 | 1.675 | 0.037 | 0.008 |
| 22 | F258-4B2-1 | 2.346 | 2.385 | 0.336 | 0.178 |
| 23 | F258-4B5-1 | 2.195 | 1.125 | 0.513 | 0.291 |
| 24 | F258-4B9-1 | 1.728 | 1.961 | 0.109 | 0.048 |
| 25 | F258-4B10-1 | 1.459 | 0.058 | 1.427 | 0.015 |
| 26 | F258-4C7-1 | 1.674 | 0.416 | 1.915 | 0.027 |
| 27 | F258-4D9-1 | 1.493 | 0.997 | 0.013 | 0.002 |
| 28 | F258-4E4-1 | 0.075 | 0.257 | 0.101 | -0.001 |
| 29 | F258-4E5-1 | 1.356 | 0.047 | 1.367 | 0.005 |
| 30 | F258-4E11-1 | 1.202 | 0.030 | 1.346 | 0.003 |
| 31 | F258-4F6-1 | 1.459 | 0.319 | 2.127 | 0.010 |
| 32 | F258-4H10-1 | 1.575 | 0.860 | 1.603 | 0.210 |
| 33 | F258-5E1-1 | 0.023 | 1.734 | 0.045 | 0.015 |
| 34 | F258-5G8-1 | 1.152 | 1.261 | 0.003 | 0.008 |
| 35 | F258-6B10-1 | 1.525 | 0.732 | 1.542 | 0.155 |
| 36 | F258-6D10-1 | 0.678 | 0.547 | 1.170 | 0.327 |
| 37 | F258-6E2-1 | 1.660 | 0.196 | 0.049 | 0.028 |
| anti-GFP | 3E6-2 | 0.069 | 0.031 | 0.027 | 0.023 |
| anti-VHH3 | F236-3F7-2 | 0.217 | 0.108 | 2.383 | 0.064 |
| anti-hlgG1 Ab | F386-1E8-1 | 0.011 | 0.009 | 0.006 | 2.315 |

**Supplemental Figure 1:**
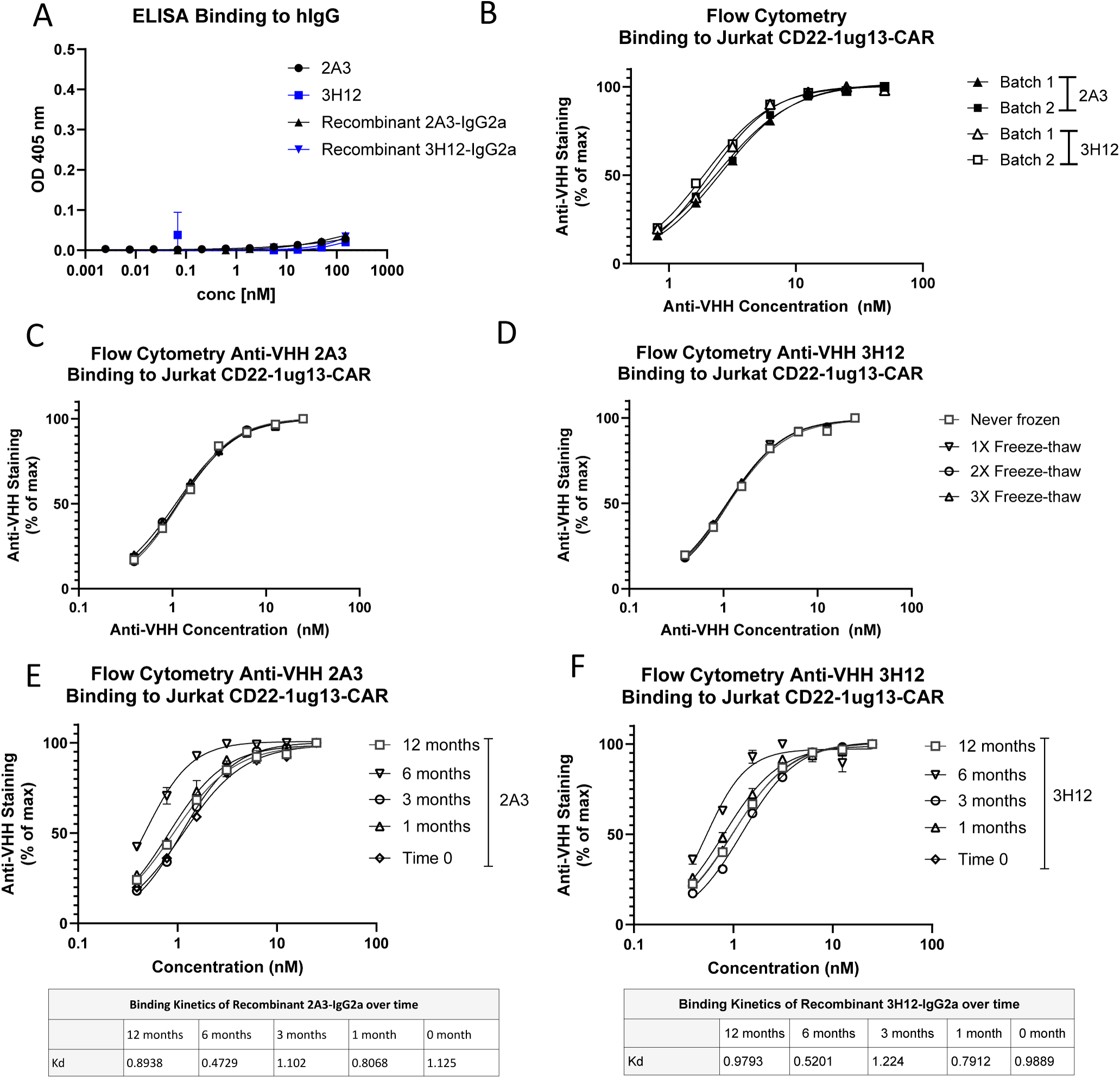
*Recombinant anti-VHH IgG2a mAbs maintain low reactivity to human IgG* Purified anti-VHH IgG1 2A3 or 3H12 mAbs, or the corresponding recombinant IgG2a antibodies were tested via ELISA against (A) total human IgG protein using anti-mouse IgG as a detection antibody. Results demonstrate that purified anti-VHH mAbs maintain high reactivity to llama VHH with low reactivity to human antibodies. Anti-VHH mAbs were then examined for production consistency using (B) assessment of binding to VHH-CAR via flow cytometry over a range of anti-VHH mAb concentration. (C-D) Stability of a anti-VHH mAb product over multiple rounds of freeze-thaw was also confirmed via flow cytometry over a range of anti-VHH concentrations. (E-F) Aliquots of anti-VHH mAbs were stored at −80°C and examined for reactivity to VHH-CAR expressing cells at various timepoints after freezing via titrated flow cytometry, confirming product stability for at least 12 months.

**Supplemental Figure 2.**
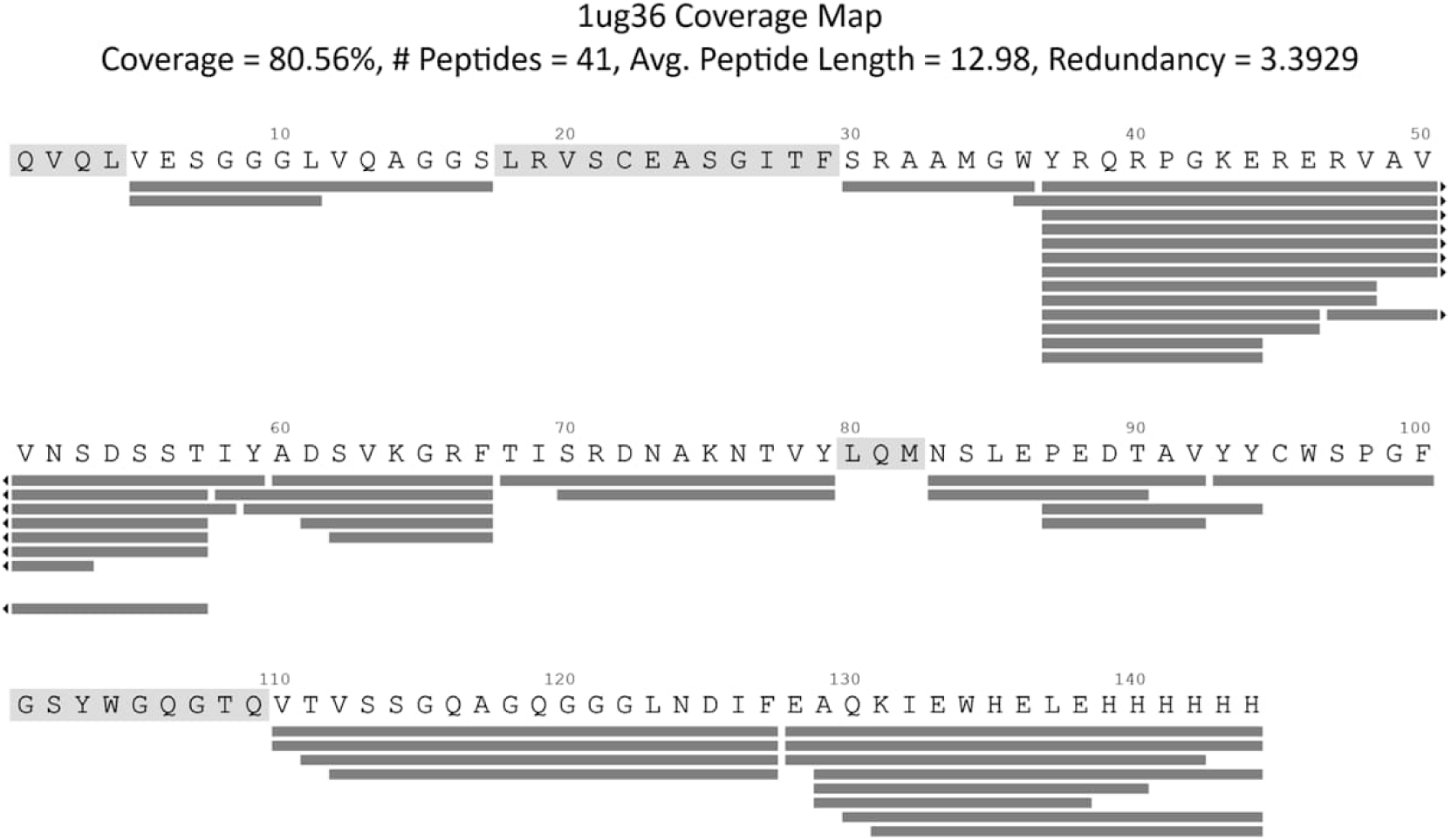
HDX-MS sequence coverage. Deuterated peptides are shown as grey rectangles below the amino acid sequence.

**Supplemental Table 2.**
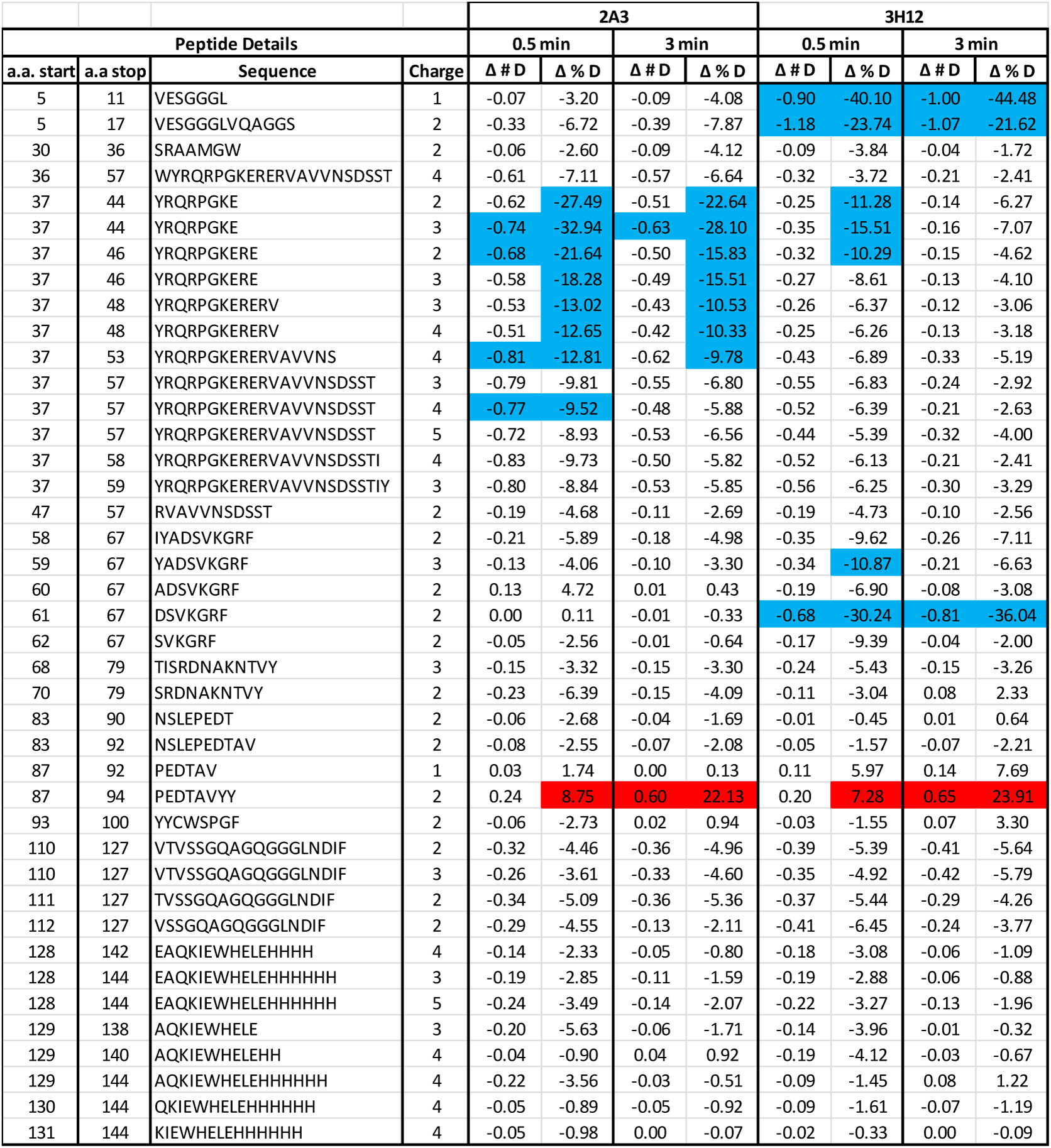
Absolute (# D) and Normalized (% D) Differential HDX-MS for each identified peptide (n = 3). Significant reductions and increases in HDX in the bound state relative to the unbound state are shown in blue and red, respectively.

**Supplemental Figure 3:**
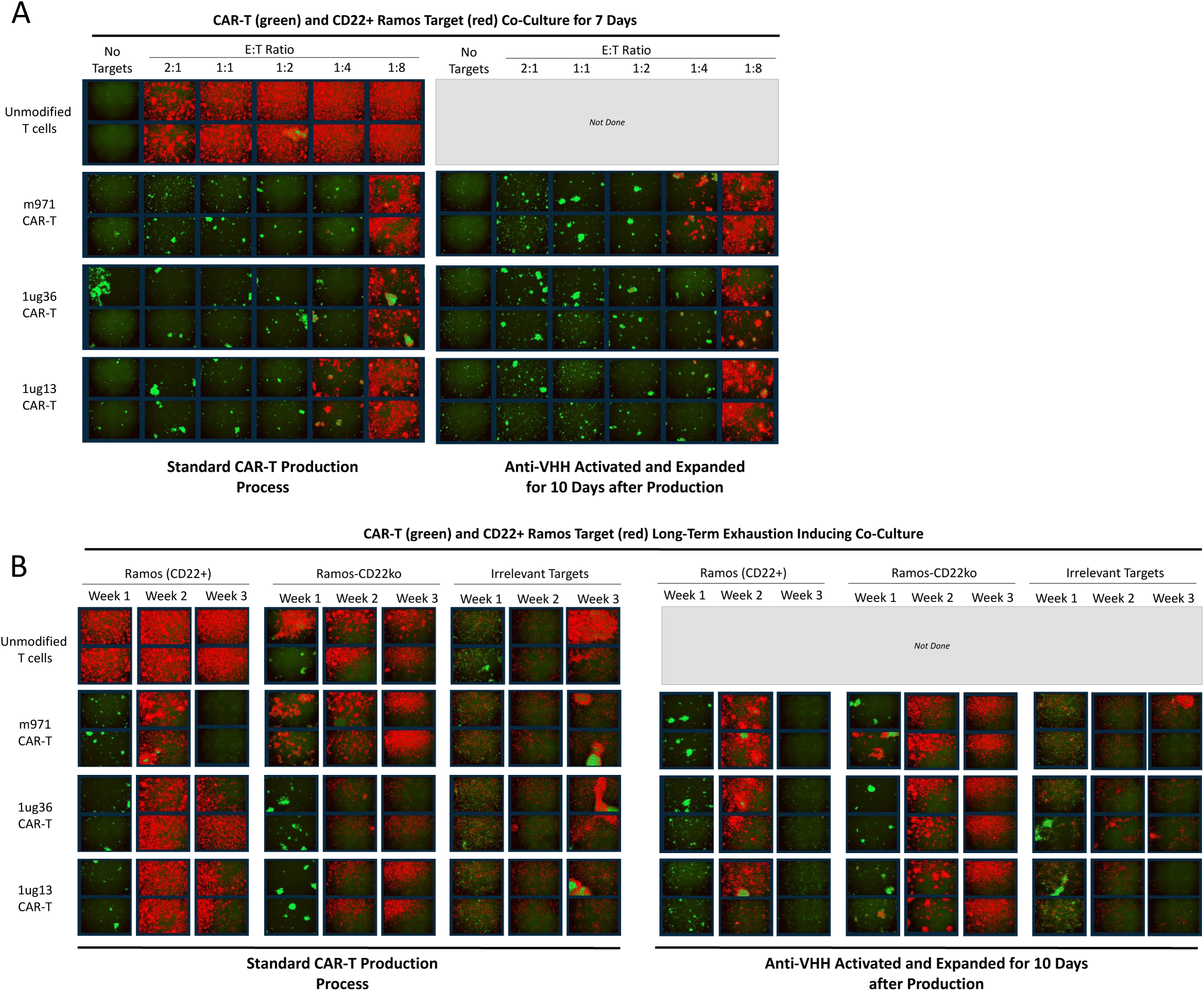
*Fluorescence images demonstrate short and long-term function of CD22 CAR-T cells with or without anti-VHH stimulation.* Cryopreserved CD22 CAR-T cells or matched untransduced T cells were activated with anti-VHH mAb coated plates, or left unactivated, and expanded for 12 days. CAR-T cells with or without activation were then placed in co-culture with red-fluorescent protein expressing target cells. (A) Co-cultures of 20 000 target cells were established with varying numbers of CAR-T cells to achieve effector to target ratios. Images show the relative number of CAR-T cells (green) and target cells (red) after 7 days of co-culture. (B) Similarly, co-cultures were monitored in a longer term exhaustion inducing assay, wherein additional targets were added twice per week, and cultures were split 1 in 5 once per week. Images show the relative number of CAR-T cells (green) and target cells (red) before cell culture splits at days 7, 14, and 23 of co-culture.

